# Structural transitions permitting ligand entry and exit in bacterial fatty acid binding proteins

**DOI:** 10.1101/2021.09.16.460654

**Authors:** Jessica M. Gullett, Maxime G. Cuypers, Christy R. Grace, Shashank Pant, Chitra Subramanian, Emad Tajkhorshid, Charles O. Rock, Stephen W. White

## Abstract

Fatty acid (FA) transfer proteins extract FA from membranes and sequester their ligand to facilitate its movement through the cytosol. While detailed views of soluble protein-FA complexes are available, how FA exchange occurs at the membrane has remained unknown. *Staphylococcus aureus* FakB1 is a prototypical bacterial FA transfer protein that binds palmitate within a narrow, buried tunnel. Here, we determine the conformational change from this closed state to an open state that engages the phospholipid bilayer. Upon membrane binding, a dynamic loop in FakB1 that covers the FA binding site disengages and folds into an amphipathic helix. This helix inserts below the phosphate plane of the bilayer to create a diffusion channel for the FA to exchange between the protein and the membrane. The structure of the bilayer-associated conformation of FakB1 has local similarities with mammalian FA binding proteins and provides a general conceptual framework for how these proteins interact with the membrane to promote lipid transfer.

## Introduction

Lipids are hydrophobic molecules with limited water solubility that are transferred between membrane organelles or to soluble enzymes by lipid transfer proteins that sequester these apolar molecules and facilitate their transfer through the cytosol (Hertzel and Bernlohr, 2000; Wong et al., 2017; Wong et al., 2019). There are five essential steps common to all lipid transfer processes (Figure 1A). First, the transfer protein ferries its enclosed cargo to the membrane. Second, it collides with the membrane bilayer surface via an electrostatic attraction or by exploiting a specific membrane ligand such as the phospholipid head group. Third, a conformational change occurs that opens the protein interior, exposes the buried lipid, and allows for its exchange with another lipid in the membrane. Fourth, the conformational change reverses and the new protein-lipid complex dissociates from the membrane. Fifth, the complex moves through the cytosol to its destination. There are detailed structures of the lipid transfer proteins in solution illustrating how they sequester lipids from solvent and allow transport through the cytosol (Storch and Corsico, 2008; Zimmerman and Veerkamp, 2002). In contrast, structures of the exchange state of transfer proteins at the membrane interface remain elusive.

**Figure 1.**
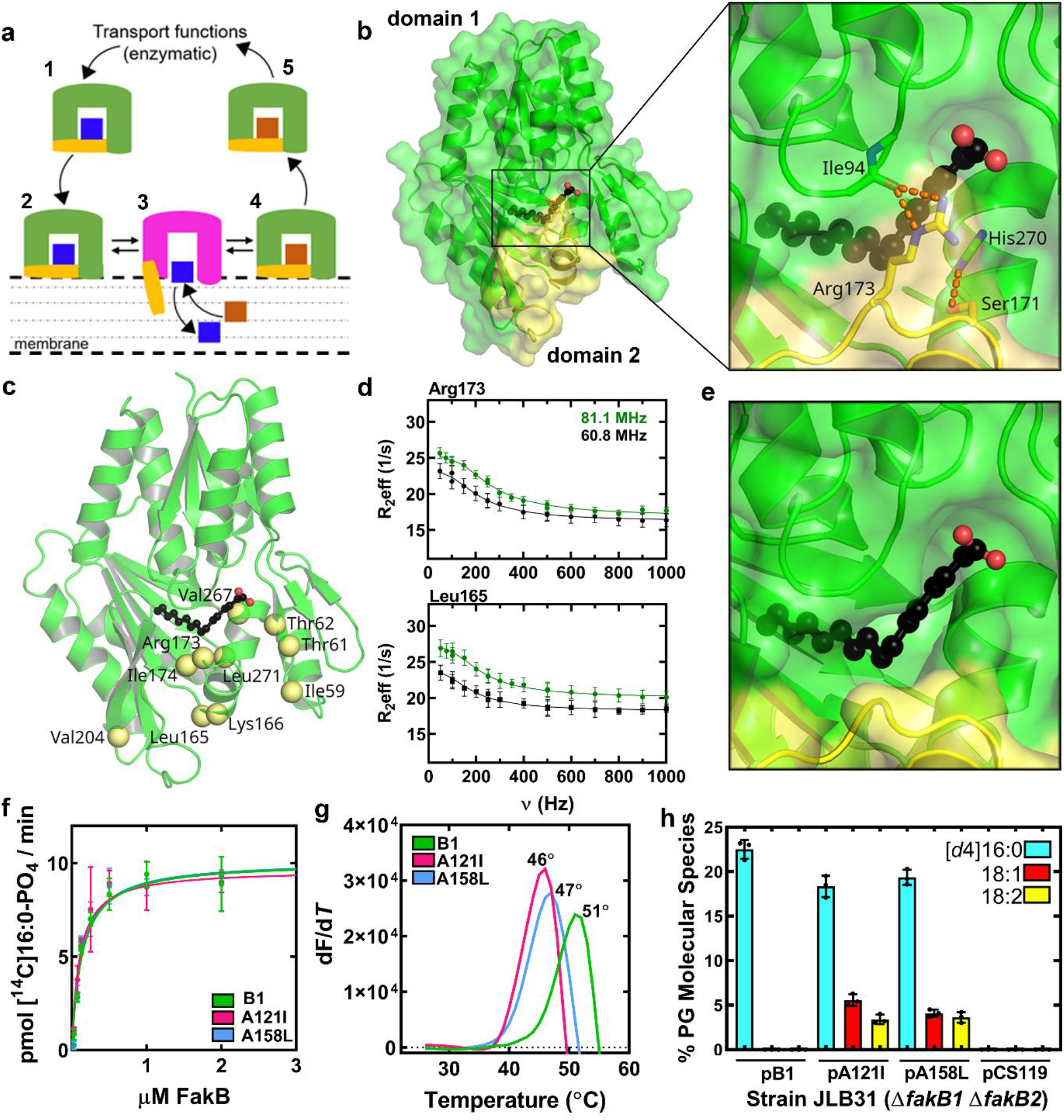
A dynamic domain in FakB1. (a) Current model for membrane lipid transfer by FA binding proteins (FABP). FABP exists in a closed conformation (1) that sequesters a FA (blue) during transfer through the cytosol, binds to the membrane (2), undergoes a conformational change (magenta) at the membrane to create a portal for FA exchange (3), and returns to the closed conformation (4) to transport a new FA (brown) (5). (b) The FakB1 domain 1 (green) and domain 2 (yellow) form a closed conformation stabilized by a hydrogen bond latch (orange dotted lines) between Arg173 and Ser171 in domain 2 and Ile94 and His270 in domain 1 that closes the domain interface over the first 8 carbons of the acyl chain of the bound FA. (c) The residues with measurable CPMG-RD exchange rates are indicated as yellow spheres mapped onto the FakB1 crystal structure. (d) The 2-state fit (lines) mapped onto the ^15^N relaxation dispersion data at two field strengths for Arg173 and Leu165. Error estimates for Reff were obtained from duplicate measurements at 100, 250, and 500 Hz as previously described{Korzhnev, 2004 #2}. (e) Closeup of the domain interface illustrating how movement of the dynamic residues could open a portal into the FA binding site. (f) The apparent K_M_s for FakB1 and mutant FakB1 binding to FakA (FakB1, 0.13 ± 0.02 µM; FakB1(A121I), 0.1 ± 0.02 µM; FakB1(A158L), 0.13 ± 0.02 µM). Mean ± SD are shown; *n* = 3 independent experiments. (g) Representative spectrum showing the thermal stabilities of FakB1, FakB1(A121I), and FakB1(A158L) using SYPRO orange. (h) Substrate selectivity of FakB1, FakB1(A121I), and FakB1(A158L) in vivo. Mean ± SD are shown; *n* = 3 independent experiments.

The mammalian fatty acid (FA) binding protein (FABP) family has a common β barrel fold formed by ten antiparallel β-strands that creates a large internal cavity to accommodate many different FA structures (Hertzel and Bernlohr, 2000; Storch and Corsico, 2008; Storch and Thumser, 2000). The amino termini of FABPs form a helix-loop-helix motif that ‘caps’ the internal cavity of the β barrel. Crystal structures show no obvious opening for an external FA to access the interior pocket, but NMR studies have revealed a dynamic helical cap region (Corsico et al., 1998; Herr et al., 1996; Storch and Corsico, 2008). Site-directed mutagenesis studies support a role for the cap helices in membrane association and the subsequent extraction of the FA (Corsico et al., 1998; Herr et al., 1996; Likic and Prendergast, 1999; Liou and Storch, 2001; Storch and Thumser, 2000; Zhang et al., 1997) while molecular dynamics (MD) simulations corroborate the dynamic nature of the two-helix motif in solution (Bakowies and van Gunsteren, 2002; Friedman et al., 2006; Guo et al., 2019; Li et al., 2015; Long et al., 2009; Matsuoka et al., 2015; Tsfadia et al., 2007). The current model posits that the cap exists in an ‘open’ state when an FABP is bound to the bilayer to allow FA exchange, but the conformation of the bilayer-associated FABP has not been described. The bacterial class of FA transfer proteins is called FakB. Like their mammalian counterparts, FakB proteins shuttle FA and acyl-PO_4_between the membrane bound and soluble enzymatic partners (Cuypers et al., 2019; Parsons et al., 2014). They are structurally distinct from mammalian FABPs and have restricted internal cavities that are tailored to bind only selected FA structures (Broussard et al., 2016; Cuypers et al., 2019; Gullett et al., 2019; Parsons et al., 2014). Like FABPs, the acyl chains in FakBs are completely enclosed within the protein interior, and the protein conformation allowing the FA to escape into the membrane is unknown.

Despite their dissimilar folds, the mammalian and bacterial FA transfer proteins face an identical topological challenge. Namely, a conformational change must occur to create a diffusion channel for the FA to travel from its position in the membrane to the interior of the transfer protein. Here, we use a combination of X-ray crystallography, NMR spectroscopy, MD simulations, site-directed mutagenesis, and functional biochemical assays to characterize the membrane-bound conformation of FakB1, a FA transfer protein from *Staphylococcus aureus* that specifically binds palmitic acid (16:0) before presenting it to FakA for phosphorylation and exchange with the membrane. FakB1 has a flexible 23- residue lid covering the FA binding tunnel, and we find here that this lid domain undergoes dynamic exchange between ‘latched’ (closed) and ‘unlatched’ (open) states in solution. We captured the crystal structure of the open FakB1 conformation by introducing point mutations into the narrow FA tunnel to destabilize the latch. The open conformation arises from the rotation of helix α8 outward to uncover the FA-binding tunnel concomitant with the reorganization of the α8-β9 flexible loop into a new helix α8’. MD simulations reveal how α8’ inserts into the phospholipid bilayer to create a diffusion path for the FA to exit into the membrane, and site-directed mutagenesis corroborates the roles of key residues in this process. These data provide a complete picture of how FA binding proteins undergo conformational changes necessary for the exchange of lipids between the protein interior and the membrane.

## Results

### A dynamic domain in FakB1

Previous FakB crystal structures show that the FA is completely buried within the protein interior with only the FA carboxylate group exposed for phosphorylation (Broussard et al., 2016; Cuypers et al., 2019; Gullett et al., 2019). The structures also suggest a potential focal point for the conformational change that must occur to release the FA. FakBs have a large domain 1 (residues range) that contains the bulk of the acyl chain binding pocket and a smaller domain 2 (Asp164-Lys186) that forms a lid over the first 8 carbons of the FA chain (Figure 1B). The two domains are fastened together over the FA by a hydrogen bond latch between the side chains of Arg173 and Ser171 on domain 2 and the backbone carbonyl of Ile94 and the Nδ1 of His270 on domain 1 (Figure 1B). Opening of the latch and the rotation of helix α8 (Leu165 to Ser171) would expose the FA and provide a portal for exchange. Here we used NMR spectroscopy to determine if there is flexibility along the domain interface of FakB1 that is not apparent from its crystal structure. The 2D [^15^N,^1^H] TROSY spectrum of [^15^N, ^13^C]FakB1 showed well dispersed resonances (Figure S1A), with all but four residues assigned at 293 K. The Cα chemical shift deviation plots correlate with the secondary structural elements observed in the FakB1 crystal structure (Figure S1B).

Carr-Purcell-Meiboom-Gill relaxation dispersion (CPMG-RD) spectra were acquired using [^2^H,^15^N,^13^C]FakB1, at two different field strengths (81.1 MHz and 60.8 MHz) at 293 K to identify and characterize mobile residues in FakB1. Ten residues exhibited ^15^N relaxation dispersion profiles, and their mapping onto the FakB1 crystal structure shows that they are either in domain 2 or adjacent to the FA and include the Arg173 latch and Leu165 at the domain junction at the amino terminus of helix α8 (Figure 1C). These two data sets exhibit close fits to a two-state exchange model using the equation of Carver and Richards (Carver and Richards, 1972) (Figure 1D). All ten residues were fit to a global 2- state model with forward (k_f_) and reverse (k_r_) exchange rate constants 12.0 ± 1.0 s^-1^ and 753.0 ± 53.0 s^-1^, respectively, yielding an exchange rate constant (k_ex_= k_f_+ k_r_) of 764.2 ± 53.0 s^-1^. A Gibbs free energy barrier of 2.41 ± 0.07 kcal/mol is calculated from the exchange rates for the FakB1 conformational change. The major (98.5%) and the minor (1.5%) conformations exchange on the millisecond time scale in solution. Removal of the ten mobile residues from the FakB1 crystal structure would create an opening along the domain interface that exposes the FA binding site (Figure 1E). Hence, the observed NMR dynamics suggest a conformational change consistent with an opening along the domain interface that would create a portal to the FA binding tunnel.

### X-ray crystallography captures an open conformation

Although NMR spectroscopy shows the Arg173 latch to be dynamic, the latch is firmly closed in the FakB1 crystal structure and a putative unlatched, open structure has never been observed. Unlike the mammalian FABPs, FakB1 has a tight binding tunnel that presents an opportunity to use site-directed mutagenesis to modulate the optimal packing of the FA into its binding site. We identified two alanine residues (121 and 158) in the tunnel and mutated them to bulkier residues to partially occlude the FA binding pocket, push the FA out of its pocket, and destabilize the fully closed FakB1 conformation. The FakB1(A121I) and FakB1(A158L) mutant proteins were fully active in FA kinase assays with the same apparent affinity for FakA (Figure 1F) and analytical ultracentrifugation verified that both form tight complexes with FakA (Table S1). However, the mutations did alter two key properties of FakB1. First, compared to the wild-type protein, the thermal stabilities of FakB1(A121I) and FakB1(A158L) were reduced by 5 °C and 4 °C, respectively (Figure 1G), showing that the mutations prevent FakB1 from adopting its most stable conformation. Second, the mutations altered the FA selectivity of FakB1 in a physiological setting. When expressed in *E. coli*, the total amount of FA incorporated by the FakB1 mutant proteins was the same as the wild-type protein, but FakB1 only supported the incorporation of palmitic acid (16:0), whereas both mutant proteins were less specific and also incorporated oleate (18:1) and linoleate (18:2) (Figure 1H). These data show that the tunnel mutations alter the stability and selectivity of FakB1, but do not impair the overall function of FakB1 in vivo or in vitro.

We determined the crystal structure of FakB1(A121I) at 2.02 Å resolution (Table 1). We observed two molecules (molA and molB) in different conformational states in the asymmetric unit. The prototypical FakB1 closed conformation was exemplified by molB, but in molA, the FakB1 domains are not in direct contact (Figure 2A). Instead, domain 2 of molA adopts a new ‘open’ state that arises from unfastening the Arg173 latch, rotating helix α8 (Leu165 to Ser171) away from the domain interface and releasing the α8-β9 loop residues (Thr175 to Lys186) from domain 1 to fold into a new 3-turn α-helix (α8’). Excluding domain 2, the open and closed structures superimpose with an RMSD of 0.356 Å showing that the conformational change is confined to residues within domain 2 (Figure 2A). Notably, the conformational change occurs precisely in the dynamic region of the protein revealed by NMR spectroscopy (Figure 1C). The surface rendering of FakB1(A121I) shows that the domain 2 conformational change exposes the ligand binding cavity and creates a portal to the FA (Figure 2B).

**Table 1.**
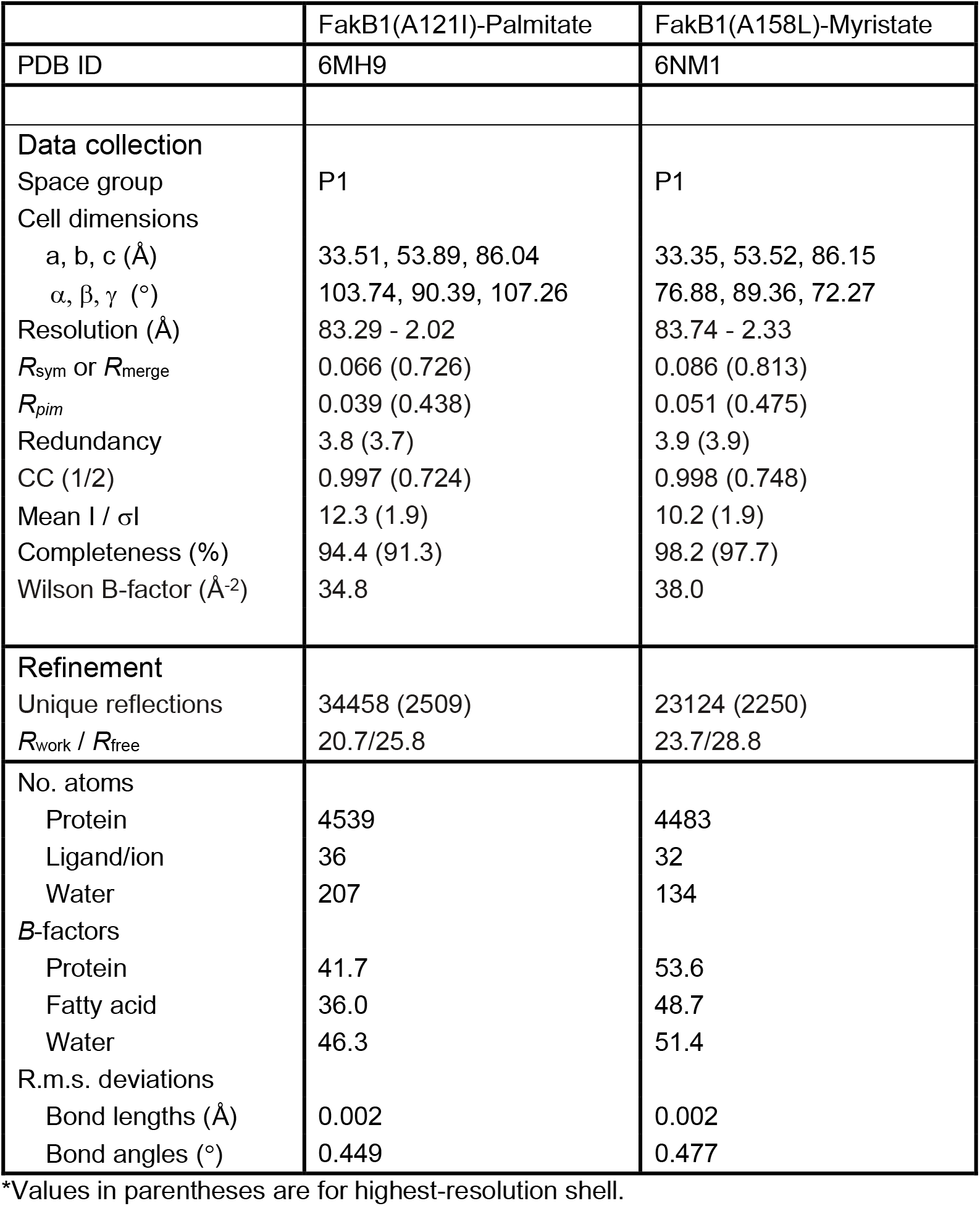
X-ray crystallography collection, refinement, and validation statistics.

**Figure 2.**
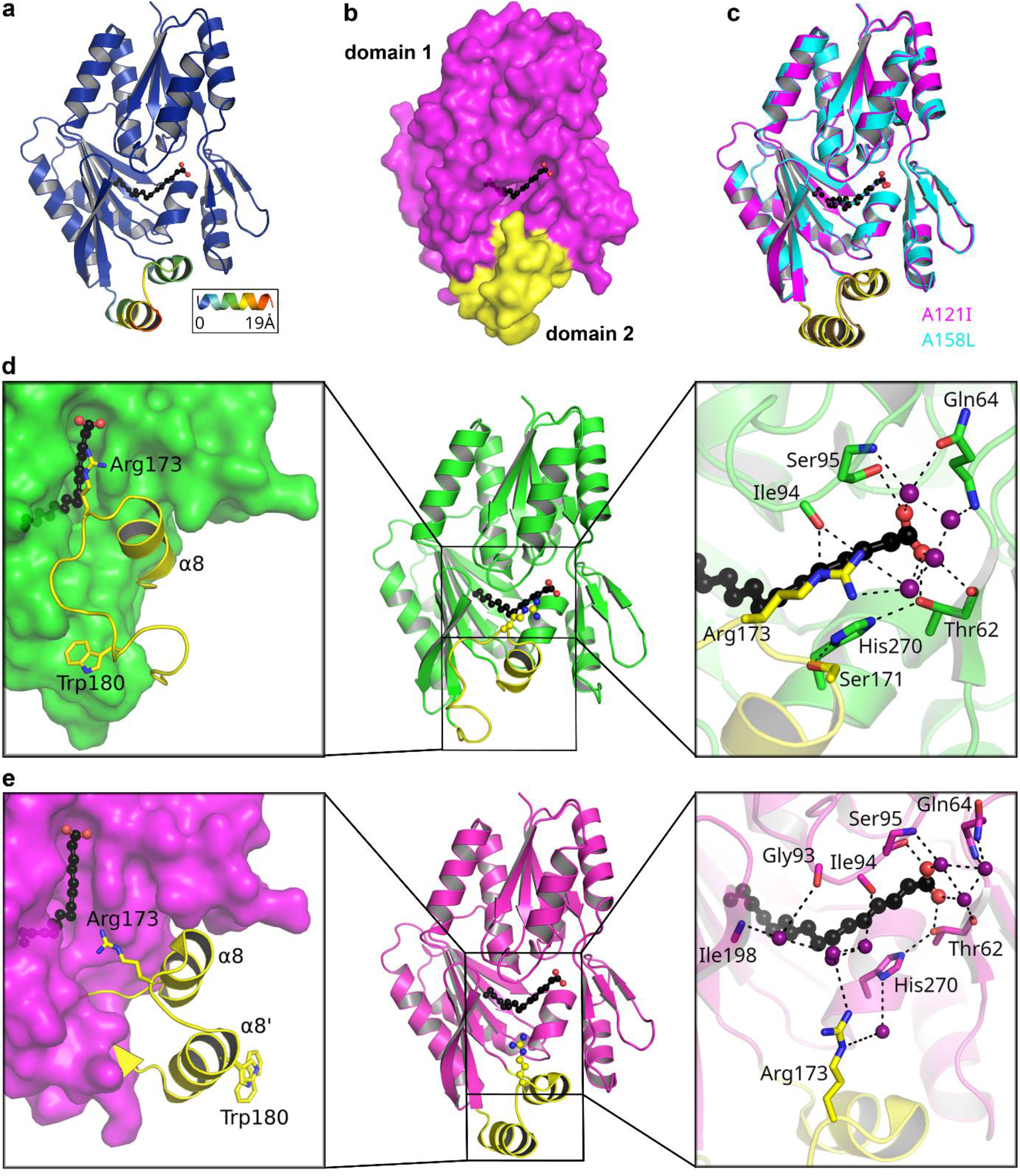
The ‘closed’ and ‘open’ states of FakB1. (a) The RMS deviations (represented by blue (low) to red (high)) in the main chain Cα positions of the FakB1(A121I) crystal structure relative to wild type FakB1 (not shown). (b) Surface rendering of open FakB1(A121I) showing how the rotation of domain 2 (yellow) away from domain 1 (magenta) creates a portal exposing the FA. (c) Overlay of the open FakB1(A121I) and open FakB1(A158L) crystal structures. (d) Domain interactions in FakB1. *Left Panel*, domain 2 (yellow) is packed tightly against domain 1 (green). Within domain 2, hydrophobic residues of the α8-β9 loop make close van der Waals interactions and the Arg173 latch at the end of helix α8 is closed. *Right Panel*, the water-mediated hydrogen bond network (black dotted lines) between Arg173 and Ser171 on domain 2 (yellow) and Ile94 and His270 on domain 1 (green), respectively. The Ser171-His270-Thr62-FA carboxylate-Ser95 hydrogen bond network and structured waters (purple spheres) at the FA carboxyl binding site are also shown. (e) Domain interactions in FakB1(A121I). *Left Panel*, movement at the base of helix α8 rotates it away and the structured loop is released from domain 1 (magenta) to form helix α8’. *Right Panel*, the latch hydrogen bond interactions are broken. Ile94, Arg173, and His270 form contacts with surrounding water molecules (purple spheres) thereby disconnecting the domains and disrupting the Ser171-His270-Thr62-FA carboxylate-Ser95 hydrogen bond network.

We also determined the 2.33 Å crystal structure of FakB1(A158L) (Table 1), which was the same in all respects to the structure of FakB1(A121I). The open FakB1(A158L) molA structure superimposes on the FakB1(A121I) molA structure with an RMSD of 0.265 Å (Figure 2C). In FakB1(A121I), the bulkier side chain displaces the middle of the aliphatic chain of the FA by 0.9 Å and its distal end by ∼1.4 Å that, in turn, shifts the tunnel residues Val162, Leu191, and Leu271 and pushes against the domain interface and the Arg173 latch. A similar process occurs in FakB1(A158L).

In the closed conformation, the hydrophobic α8-β9 loop containing Trp180 is tightly packed against the hydrophobic surface of domain 1 and the Arg173 latch is closed (Figure 2D, left panel). There is also a hydrogen bond network consisting of Ser171, His270, Thr62, Ser95 and the FA carboxylate that appears to balance the negative charge on the FA (Figure 2D, right panel). In the open conformation, the α8-β9 loop separates from domain 1 and folds into helix α8’, and the Arg173 latch is unfastened with helix α8 rotated away from the domain interface (Figure 2E, left panel). The newly formed helix α8’ has a distinct hydrophobic surface consisting of Ala177, Trp180, Val181, Leu184, and Leu185. The low dielectric constant within the FakB1(A121I) crystal lattice coupled with the location of Phe38 and Ile44 on the neighboring molecule that is only available to molA create a favorable environment for the formation of helix α8’ (Figure S2A). The opening of the Arg173 latch and outward movement of α8 disrupt the hydrogen bond network surrounding the FA carboxylate by breaking the Ser171-His270 hydrogen bond and exposing His270 to solvent (Figure 2E, right panel).

### FakB1(A121I) dynamics

FakB1(A121I) was analyzed by NMR to determine how the mutation impacts protein dynamics. The 2D [^15^N,^1^H] TROSY spectrum of FakB1(A121I) was similar to that of the wild type protein (Figure S2B) with the largest chemical shift differences localized to the area adjacent to Ala121 (Figure. S2C). CPMG-RD data for [^2^H,^15^N,^13^C]FakB1(A121I) collected at two field strengths at 293 K revealed 12 residues with ^15^N relaxation dispersion profiles that mapped to the same locations as in FakB1 (Figure S2D). The data sets for Arg173 and Leu165 reveal the flexibility of the latch and base of helix α8 in FakB1(A121I) (Figure S2E). The global exchange rate was k_ex_= 757.3 ± 42.0 s^-1^, with forward (k_f_) and reverse (k_r_) exchange rate constants, 36.0 ± 3.0 s^-1^ and 722.0 ± 42.0 s^-1^ respectively. The population of the minor species (p_minor_) increased from 1.5% in FakB1 to 5% in FakB1(A121I). The lower Gibbs free energy barrier (ΔG = 1.75 ± 0.02 kcal/mol) suggests that FakB1(A121I) is less stable than FakB1 consistent with its lower melting temperature (Figure 1G).

Although the NMR experiments show that domain 2 is flexible compared to the rigid domain 1, they do not establish the major and minor species in solution. We therefore analyzed specific ^N^H-^N^H NOEs in the ^15^N-resolved NOESY spectrum. The high quality of the NOEs was confirmed by the strong ^N^H^i^-^N^H^i+1^ and ^N^H^i^-^N^H^i+2^ NOEs for helix α4 in both FakB1 and FakB1(A121I) (Figure S3). The conformation of the domain 2 α8-β9 loop in FakB1 is revealed by the Thr175-^N^H to Ile198-^N^H NOE and the reverse Ile198-^N^H to Thr175-^N^H NOE that show Thr175 on the α8-β9 loop is in close proximity to Ile198 on β10 of domain 1 (Figure 3A). The same NOE connections between the sidechain aromatic protons of Trp180 to Glu202-^N^H and Lys203-^N^H observed in FakB1 (Figure 3A) are also detected in FakB1(A121I) (Figure S4A). The NOE-based distances for FakB1 and FakB1(A121I) are consistent with the distances in the crystal structure of the closed conformation and are inconsistent with the distances in the open conformation, i.e., formation of helix α8’ in solution.

**Figure 3.**
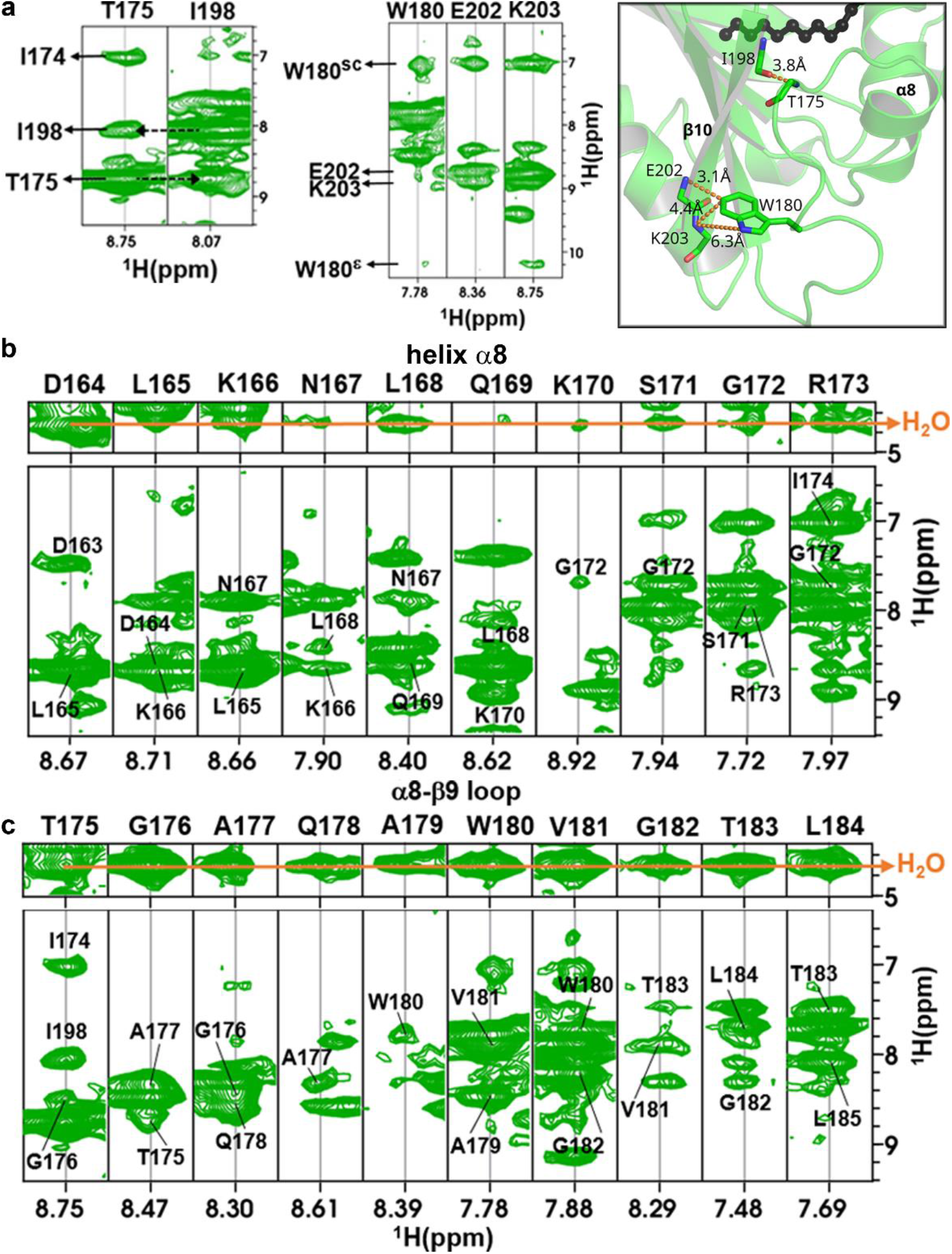
Solution conformation of FakB1. (a) NOE contacts between the α8-β9 loop of domain 2 and residues in domain 1 are consistent with the closed conformation crystal structure. The NOE interactions (dotted orange lines) and computed distances from the NMR data are shown (sc, side chain protons; ε, amide protons). (b) Sequential NOEs confirm the existence of helix α8 in solution. The orange line indicates exchange cross peaks with water. (c) The water cross peaks from Thr175-Leu184 (orange line) are indicative of a structured α8-β9 loop, but the sequential NOEs centered on Trp180-Val181 suggest partial helical character in this region.

Although the sequential NOEs from Asp164 to Arg173 in FakB1 are consistent with the existence of helix α8 in solution, stronger exchange cross peaks with water in Asp164- Lys166 and Ser171-Arg173 point to helix α8 being slightly unraveled at both ends (Figure 3B). In addition, although the Thr175-Leu184 α8-β9 loop residues have strong water cross peaks consistent with the loop conformation in the X-ray structure (Figure 3C), the sequential NOEs across this region suggest that the loop has some helical characteristics centered on Trp180-Val181. The crystal structure of FakB1 reveals that there is indeed a single helical turn between residues Gly176 and Val181 (Figure 2D, left panel). These observations are confirmed by the analysis of the NMR spectra of helix α8 (Figure S4B) and the α8-β9 loop (Figure S4C) of FabB1(A121I). Taken together, the NOE data are consistent with the closed FakB1 conformation as the major state in solution.

### FakB1 membrane interactions

FakB1 must bind to the membrane and create a path for the FA to travel from its location in the phospholipid bilayer (Pashkovskaya et al., 2018) and into the protein binding site, and vice versa. Our structural and dynamic analyses strongly suggest that an open conformation of FakB1 mediates this process, and we tested this hypothesis by investigating the binding of FakB1 to charged phospholipids.

Phosphatidylglycerol (PG) is the major phospholipid in the inner bilayer of the *S. aureus* membrane and a PG liposome pull-down experiment reveals that FakB1 has an affinity for PG, but not for phosphatidylcholine (PC) liposomes (Figure 4A). We then determined the affinities of FakB1 and FakB1(A121I) for PC:PG (50/50) vesicles using PC vesicles as a negative control in surface plasmon resonance (SPR) assays. We detected a 15-20 μM affinity of both FakB1 and FakB1(A121I) for PG-containing vesicles (Figure 4B), which is similar to its micromolar affinity for FakA and consistent with its transport function.

**Figure 4.**
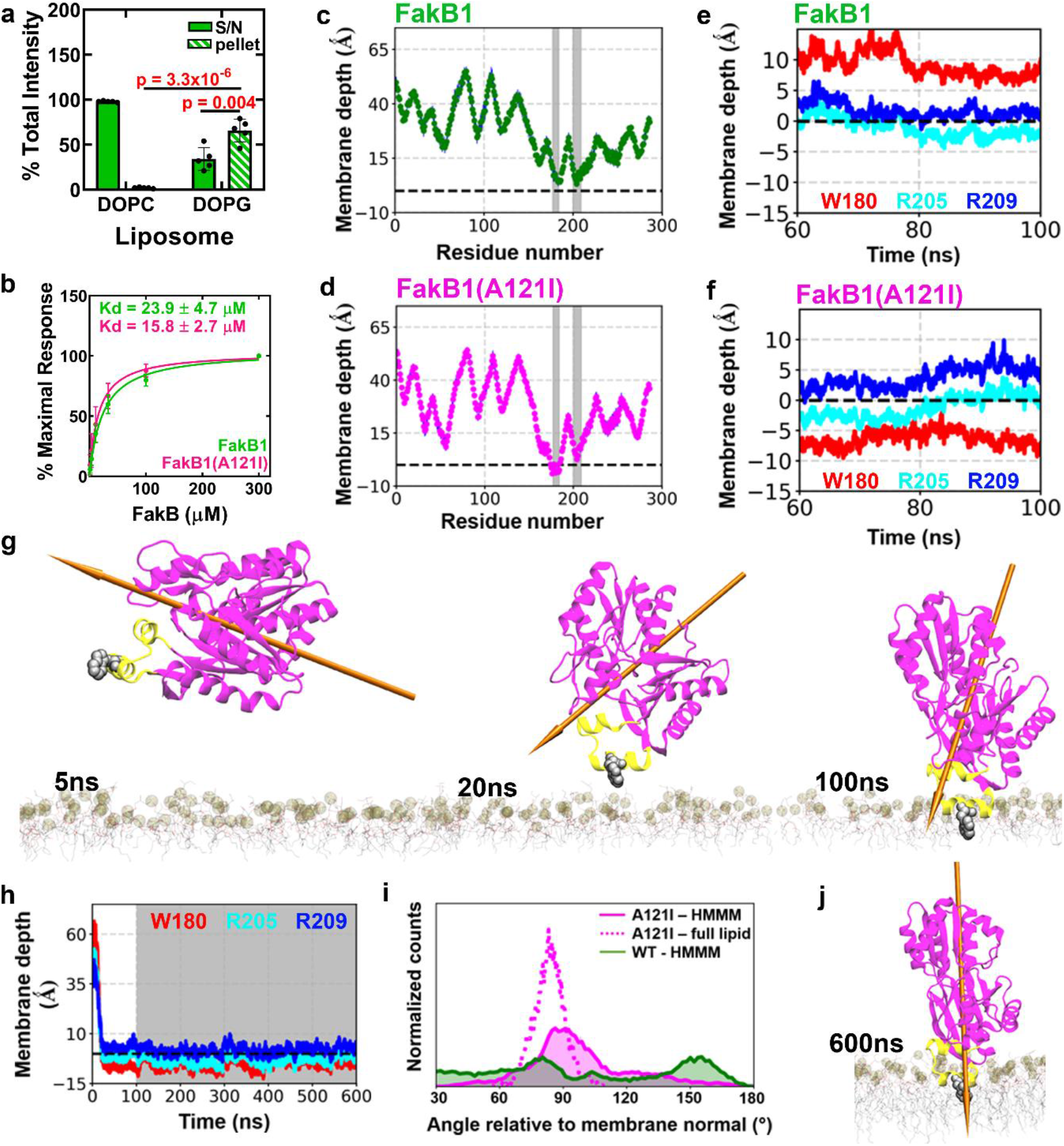
The open conformation of FakB1 inserts into the PG bilayer. (a) FakB1 association with PG, but not PC, liposomes was assessed using a liposome pull-down experiment. Mean ± SD is shown, *n* = 5 independent experiments. Two-tailed Student’s t tests were used to evaluate statistical significance. (b) SPR analysis of FakB1 and FakB1(A121I) showing similar binding kinetics using PG:PC vesicles (50/50). Kd values were calculated from 4 independent experiments for each protein. Best fit values ± SE are shown. (c) and (d) Representative ensemble-averaged location of FakB1 and FakB1(A121I) Cα residues along the membrane normal (z) axis of the PG bilayer calculated during last 50 ns of the 10 HMMM membrane binding simulations. The phosphate layer of the lipid bilayer is set as a reference at z=0 shown with a dashed line. Gray bars represent residues adjacent to or penetrating the bilayer. (e) and (f) Locations of the center of masses (COM) of Trp180, Arg205, and Arg209 sidechains along the membrane normal (z axis) of the PG bilayer (dotted black line) plotted during the last 40 ns of FakB1 and FakB1(A121I) HMMM membrane binding simulations. (g) Snapshots of the HMMM simulation of FakB1(A121I) at the indicated times shows evolution of the binding event and the orientation of the FakB1(A121I) dipole moment (arrow). (h) Time evolution of (COM) distances for Trp180 (red), Arg205 (cyan), and Arg209 (blue) with respect to the membrane normal (dotted black line) (z=0). An HMMM membrane binding simulation was performed for the first 100 ns (white background) capturing FakB1(A121I) binding to the membrane within the first 20 ns, followed by a ‘full-lipid’ simulation for additional 500 ns (gray background) showing the stability of the bound configuration. (i) FakB1(A121I) binds with helix α8’ positioned at a ∼90° angle relative to the membrane normal (z axis) that is most pronounced in the ‘full-lipid’ simulation. (j) FakB1(A121I) protein dipole moment is aligned with the z-axis of the membrane bilayer in the ‘full-lipid’ simulation.

We developed a linear, coupled biochemical assay using FakA to measure the FakB1 FA exchange/transport activity. The phosphorylation of [^14^C]16:0 in a PG liposome as substrate yielded a transfer rate of 1.4 ± 0.2 nM/min/μg, whereas the transfer rate was two orders of magnitude slower (10.7 ± 0.6 pM/min/μg) when the [^14^C]16:0 was presented as a bovine serum albumin (BSA) complex (Figure S5A). These data indicate that FakB1 is designed to efficiently extract FA from PG bilayers rather than exchange with free or protein-solubilized FA.

### Molecular dynamics (MD) simulations of FakB1 binding to membrane and FA release

MD simulations employing the HMMM (highly mobile membrane mimetic) model of the membrane (Baylon et al., 2013; Blanchard et al., 2014; Ohkubo et al., 2012; Pant and Tajkhorshid, 2020; Pogorelov et al., 2014; Soubias et al., 2020) were used to study the interaction of FA-bound FakB1 with the membrane. HMMM uses short-tailed lipids to significantly enhance the mobility of lipids, thereby facilitating spontaneous protein insertion into a bilayer. Ten replicates of each MD simulation condition were obtained by initially placing the protein 10 Å away from the bilayer surface in a random orientation. Simulations of the closed conformation with PG show that FakB1 consistently engages the membrane and orients its electropositive surface, centered on Arg205, toward the electronegative PG bilayer (Figure 4C; Figure S5B). Although this membrane association is consistently observed in multiple simulation replicates, it is not intimate, as evidenced, e.g., by the ensemble-averaged Cα positions of Trp180, Arg205, and Arg209, being still above the phosphate plane by 6.6 ± 1.9 Å, 5.5 ± 0.9 Å, and 9.0 ± 0.8 Å, respectively (Figure 4C; Figure S5B). In contrast, simulations with the FakB1(A121I) open conformation consistently show residues 177-184 (α8’) penetrating the bilayer significantly. Although the Cα carbons of Arg205 and Arg209 are in similar locations relative to the phosphate plane in the open and closed conformation simulations (5.5 ± 1.1 Å and 10.4 ± 1.2 Å, respectively, in the open conformation), the Cα carbon of Trp180 has flipped from 6.6 Å above the phosphate plane in the closed conformation to 3.1 ± 1.2 Å below the phosphate plane in the open conformation of the protein (Figure 4D; Figure S5C). Simulations using PC-only bilayers did not show any membrane association for the open conformation (Figure S5D).

Given the large size of the side chains in Arg and Trp, the position of Cα atoms may not fully capture the penetration of these residues into the membrane. To determine the depth of membrane insertion, we monitored the time evolution for the center of masses of the side chain atoms of Trp180, Arg205, and Arg209 in membrane-binding simulations (Figure S6A-S6C). In the closed state, Trp180 and Arg209 side chains are 9.0 ± 2.4 Å and 2.9 ± 1 Å, respectively, above the phosphate plane of the bilayer, whereas the Arg205 sidechain reaches the phosphate layer (−0.13 ± 0.9 Å) (Figure 4E). In the open state, Arg209 sidechain remained 3.4 ± 1.6 Å above the phosphate plane and Arg205 continues to engage the phosphate layer (1.6 ± 1.6 Å). However, the Trp180 sidechain is deeply buried in the bilayer at a depth 6.5 ± 1.2 Å below the phosphate layer (Figure 4F). These data show that Arg205 and the surrounding electropositive surface play a key role in initially orienting the protein and engaging the bilayer in both the closed and open conformations. However, the final location of Trp180 in the membrane-bound complexes is very different with its center of mass shifting ∼12 Å in the transition between the closed and open conformations.

Snapshots of FakB1(A121I) during a typical HMMM membrane binding simulation show that it orients properly with respect to the membrane and engages it within 20 ns; subsequently, the protein forms a stably membrane-associated complex with the PG bilayer between 20-100 ns (Figure 4G; Figure S5B). The HMMM approach was validated by performing a ‘full-lipid’ MD simulation of the FakB1(A121I) open conformation that extended to 500 ns beyond the HMMM membrane binding simulations. This full-lipid simulation maintained a stable association of the open conformation with the PG bilayer with the same mode and depth of membrane insertion, as measured by monitoring the locations for the center of masses for Trp180, Arg205, and Arg209 (Figure 4H). There is a more pronounced orientational preference for membrane docking of helix α8’ in the full- lipid simulation, with helix α8’ positioned parallel to the phosphate plane and the protein dipole moment aligned with the membrane normal (z axis) (Figure 4I – 4J). Thus, both the HMMM and full tail simulations capture stable membrane-bound for the open conformation and provide similar insights into interaction mode of the protein with the PG bilayers.

### Validation of the MD model

The MD simulations strongly suggest that Trp180 and Arg205 are key residues in membrane binding of the protein. We therefore prepared FakB1(R205A), FakB1(R205E) and FakB1(W180E) mutants to experimentally test this conclusion. FakB1 was not substantially destabilized by these mutations (Figure S6D) but SPR experiments showed that all three mutant proteins were defective in binding to PG vesicles (Figure 5A). Furthermore, the mutants showed reduced FA exchange activity compared to wild-type FakB1 when [^14^C]16:0 was presented in a PG liposome (Figure 5C). In contrast, when [^14^C]16:0 was delivered in BSA, the wild-type and mutant proteins were all active although the FakB1(W180E) exchange rate is higher in this assay (Figure 5B). Thus, all mutant proteins were able to mediate catalysis using BAS-solubilized FA but were defective in acquiring FA from a liposome. These data strongly support the roles of these residues in membrane binding as suggested from the MD simulations.

**Figure 5.**
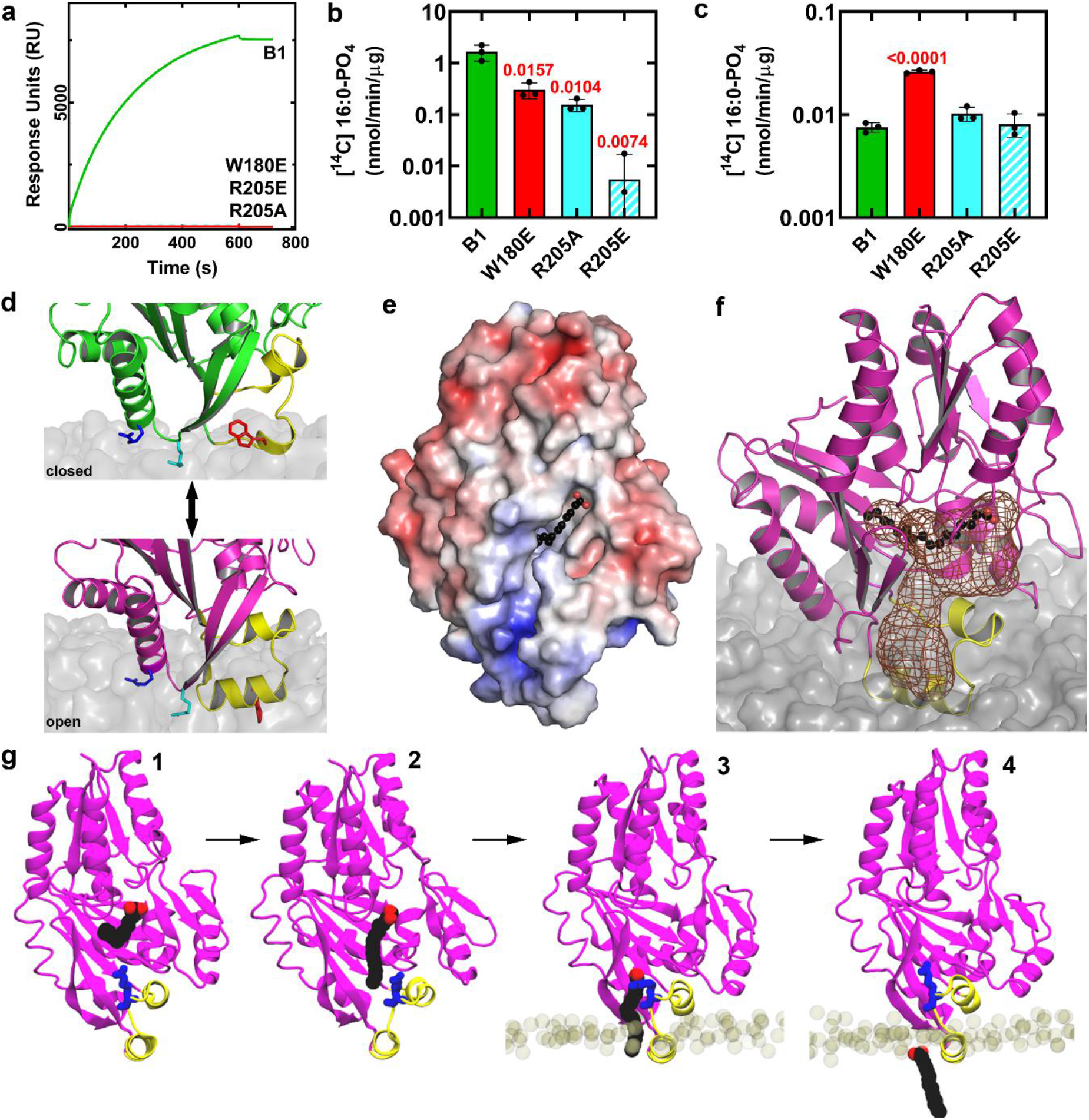
Role of Arg205 and helix α8’ in FakB1 membrane interactions. (a) Representative SPR sensogram of FakB1, FakB1(R205E), FakB1(W180E), and FakB1(R205A) binding to PC:PG (50/50) vesicles. (b) FakB1 exchange assay using [^14^C]16:0 presented in PG liposomes. (c) FakB1 FA exchange assay using [^14^C]16:0 presented as a BSA-16:0 complex. For b, c, statistical significance was determined using a two-tailed Student’s t test and comparing mutant protein values to wild-type FakB1. Mean values ± SD are shown; *n* = 3 independent experiments. (d) Closeup of the reversable conformational changes allowing FakB1 association and disengagement from a PG bilayer. *Top*, Trp180 (red) of the closed conformation is packed against the protein, the Arg205 (cyan) sidechain interacts with the phosphate layer and the Arg209 (blue) sidechain lies along the bilayer surface. *Bottom*, the open conformational change inserts helix α8’ with Trp180 pointing into the center of the bilayer and helix α8’ oriented parallel to and below the phosphate plane of the bilayer. Arg205 interacts with the phosphate groups of the PG bilayer, and Arg209 associates with the bilayer surface. (e) Electrostatic charge (red, negative to blue, positive) surface rendering of the open FakB1(A121I) crystal structure shows that a surface groove is created from the FA binding site to the electropositive end of the protein. (f) Mesh (chocolate) shows the groove becomes a diffusion channel when it is embedded in the bilayer that extends from the FA binding site in the protein interior to the phosphate plane of the bilayer. (g) Snapshots from the FakB1(A121I) **Movie S1** showing the steps in FA release into the bilayer: (1) The beginning of the simulation with the FA in its binding site. (2) The FA tail moves out of its deep pocket and into the space created by the outward rotation of helix α8 and the FA carboxyl is released from the hydrogen bond network. (3) The FA slides down the tunnel interacting with Arg173 (blue). (4) The FA diffuses into the bilayer.

The MD simulations posit a model where the electropositive surface of FakB1 surrounding Arg205 initially orients the closed conformation with respect to the membrane surface with Arg205 poised to interact with the phosphate layer (Figure 5D). The opening of the Arg173 latch and formation of helix α8’ triggers membrane penetration of the open conformation and the stabilization of helix α8’ within the bilayer (Figure 5D). Analysis of the open FakB1(A121I) crystal structure reveals a groove created by the conformational change that forms a channel extending from the FA binding site to the electropositive end of the protein (Figure 5E). In the simulations, this channel becomes partially buried when the open conformation is bound to a PG bilayer to create a hydrophobic diffusion path for the FA to travel from the protein interior to the phosphate plane of the bilayer (Figure 5F). The details of this process are revealed in one of the FakB1(A121I) MD simulations in which the FA disengages from the binding pocket, travels down the diffusion path and is fully released into the bilayer (Movie S1). This process consists of 4 steps (Figure 5G). The FA tail that is initially bound in the deep FakB1 hydrophobic pocket first moves into the portal created by the rotation of helix α8 in the open conformation. The FA carboxyl then disengages from the weakened hydrogen bond network and allows sliding of the whole FA down the tunnel along the hydrophobic surface of domain 1 that is buried by the α8-β9 loop in the closed conformation. The FA journey is interrupted for a short time by its interaction with Arg173, but then it is released to insert into the membrane where it can diffuse within the plane of the bilayer. A separate, initially membrane-associated FA can then enter FakB1 by the reverse process. The rapid conformational change coupled with the μM affinity of FakBs for PG bilayers allow fast FA exchange between the FA transfer protein and the membrane.

## Discussion

### The FakB FA exchange cycle

This study leads to a complete model for the FakB FA exchange cycle by revealing the molecular details of the membrane-bound transfer protein conformation (Figure 6). A key feature of the closed FakB conformation is the positively charged patch surrounding Arg205 that orients the protein with respect to the negatively charged PG bilayer and associates with the bilayer surface through the interaction of the Arg205 sidechain with the phosphate groups of PG. The surface-associated FakB converts to the open conformation that tightly docks to the membrane by the insertion of the new helix α8’ below the phosphate plane of the bilayer. This conformational change creates a diffusion path for the bound acyl-PO_4_/FA to release into the bilayer and for FakB to accept another FA. Thus, the transition to the open conformation in the bacterial FA binding proteins achieves four essential mechanistic goals: it opens a portal to the FA binding pocket, disrupts the hydrogen bonding network that holds the FA carboxylate in place to facilitate its release, forms a new α-helix that inserts into the bilayer, and creates a diffusion channel between the protein interior and the phosphate plane of the bilayer where FA resides. The loaded FakB protein then returns to the closed conformation, dissociates from the membrane, and delivers its cargo to FakA for another round of phosphorylation. FakB(acyl-PO_4_) also may release its cargo by interaction with cytosolic PlsX to transfer the acyl-PO_4_to acyl carrier protein or potentially with PlsY to transfer of acyl-PO_4_to glycerol-3-phosphate and initiate membrane phospholipid biogenesis (Cuypers et al., 2019).

**Figure 6.**
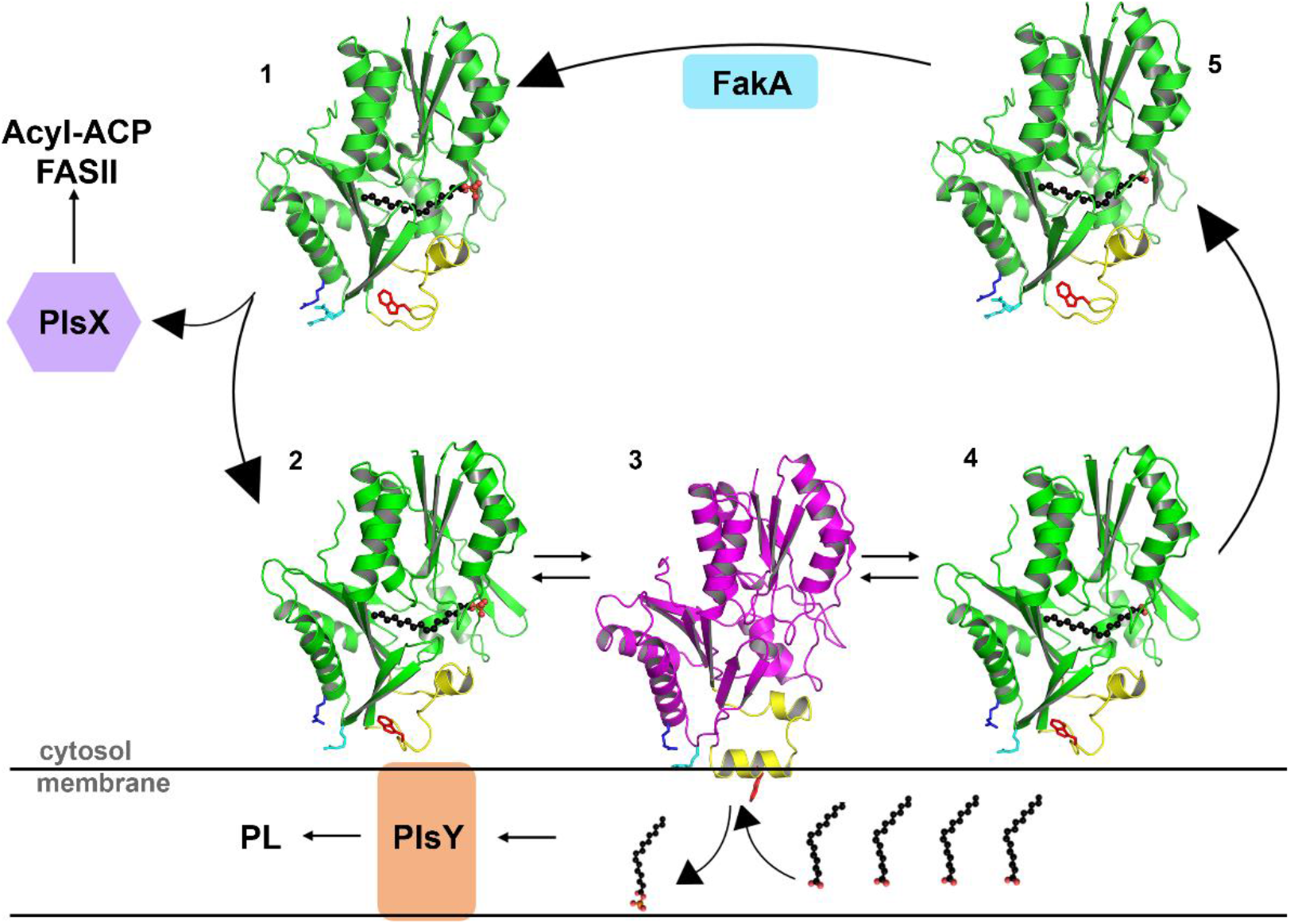
Conformational cycling model for FA exchange at the bilayer interface. (1) Phosphorylation of the FA carboxylate bound to the closed FakB1 conformation in the cytosol. (2) The electropositive surface of FakB1 is attracted to the electronegative PG membrane surface; R205 (cyan), R209 (blue), and W180 (red) are shown. (3) Domain 2 (yellow) transitions to the open conformation and helix α8’ inserts below and parallel to the phosphate plane of the bilayer with Trp180 (red) fully inserted. This FakB1-PG interaction creates a diffusion channel for FA allowing the bound acyl-PO_4_ to exchange with another FA in the bilayer. (4) Domain 2 transitions back to the closed conformation and FakB1 separates from the membrane (5) to carry its cargo to FakA to initiate another cycle of FA transfer/exchange.

### Domain 2 is metastable and switches conformations

Domain 2 of FakB that undergoes the conformational change has dynamic properties that are not obvious from the crystal structure but were revealed by the NMR analyses. These dynamic properties render domain 2 metastable and poised to switch conformations when it encounters a lipid bilayer that stabilizes the newly formed hydrophobic helix α8’. In solution, FakB oscillates between two states, a major state that corresponds to the closed structure and a minor, unlatched state that represents the first step in the transition to the open structure. The FakB1(A121I) and FakB1(A158L) crystal structures capture the fully open conformation due to the mutations that prevent full closure of the FA binding pocket and the crystal packing architecture that creates a surrogate hydrophobic environment favoring the formation of helix α8’. Arg173 and Arg205 are completely conserved in the FakB protein family and are essential for multiple FakB functions. Arg205 is at the center of the positively charge surface patch that forms the initial membrane encounter site in the MD simulations and its side chain maintains contact with the bilayer phosphate groups in both the closed and open conformations. Arg205 is also the key residue responsible for the binding of FakB to FakA (Broussard et al., 2016). Arg173 is required in the FA kinase catalytic mechanism (Broussard et al., 2016) precluding the biochemical evaluation of it role in FA exchange by site-directed mutagenesis. However, MD simulations suggest that Arg173 orients the FA with respect to the entrance to the diffusion channel and may escort the FA to and from the bilayer. Arg173 is positioned to act as a ‘lure’ dangling just above helix α8’ to capture a FA located at the phosphate zone of the bilayer and facilitate its movement into the protein’s binding pocket as FakB assumes the closed conformation and separates from the membrane. Trp180 is a sentinel hydrophobic residue whose significant conformational change and deep insertion into the membrane, captured by MD simulations, illustrates how the open conformation is anchored into the membrane.

### Similarities to mammalian FABPs

The 2-helix membrane insertion motif in the FakB1 open conformation has intriguing parallels with the 2-helix domain in mammalian FABPs (Storch and Corsico, 2008; Storch and McDermott, 2009). X-ray structures show that the prototypical FABP has two domains: a large β-barrel bottle that holds the FA attached by a hinge to a helix-turn-helix domain that caps the bottle. Although none of the solution or crystal structures have captured an open FABP conformation, it is hypothesized that the FA enters and exits the bottle via the rotation of the 2-helix cap motif (Corsico et al., 1998; Hodsdon and Cistola, 1997a, b; Sacchettini et al., 1988). NMR dynamics measurements reveal that an opening between the β-sheets may function as an alternate portal (Guo et al., 2019; Xiao et al., 2016); however, simply providing an opening for the FA is not sufficient to deliver the FA to the lipid bilayer. Previous MD simulations with FABP4 and a membrane surface (Mihajlovic and Lazaridis, 2007; Xiao et al., 2020) show that the 2-helix bundle associates with the membrane via electrostatic interactions, as suggested by extensive mutagenesis and crosslinking experiments (Corsico et al., 1998; Falomir-Lockhart et al., 2006; Herr et al., 1996), but the alternate portal between the β sheets is pointing away from the bilayer. The membrane penetrating 2-helix bundle in the open FakB1 conformation overlays perfectly with the 2-helix bundle in FABPs (Figure S6e) suggesting that the FABP4 helices may function in a similar manner to the FakB1 2-helix bundle by rotating away from the bottle opening and penetrating into the membrane to create a diffusion path for the FA. Thus, the structurally distinct FakB1 and FABP4 may both utilize a 2-helix bundle to interact with the bilayer to solve the identical topological problem faced by FA binding proteins.

## Supplementary Information

Supplementary information is provided.

## Acknowledgements

We thank Karen Miller for protein purification, Pamela Jackson for molecular biology, Matthew Frank for mass spectrometry analysis, Amanda Nourse for sedimentation analyses and Brett Waddell for SPR analyses. This work was supported by National Institutes of Health grants GM034496 (C.O.R.), and P41-GM104601 and R01-GM123455 (E.T.), Cancer Center Support Grant CA21765 and the American Lebanese Syrian Associated Charities. The diffraction data were collected at Southeast Regional Collaborative Access Team (SERCAT) beam lines 22-ID and 22-BM at the Advanced Photon Source, Argonne National Laboratory, and we thank SERCAT staff for their assistance. Use of the Advanced Photon Source is supported by the U.S. Department of Energy under Contract No. W-31-109-Eng-38. Supporting SERCAT institutions may be found at www.ser-cat.org/members.html. The simulations reported in this study are supported by XSEDE allocation (grant MCA06N060 to E.T.), Microsoft Azure, and Blue Waters at the National Supercomputing Application (NCSA) at the University of Illinois at Urbana-Champaign. The content is solely the responsibility of the authors and does not necessarily represent the official views of the National Institutes of Health.

## Author Contributions

J.M.G., M.G.C., C.R.G., S.P., C.S., E.T., C.O.R. and S.W.W. designed the study. J.M.G., C.S. and C.O.R. performed and analyzed the biochemical experiments. M.G.C. and S.W.W. determined and interpreted the crystal structures. C.R.G. performed and analyzed the NMR studies. S.P. and E.T. performed and analyzed the molecular dynamics simulations. All authors contributed to and approved the manuscript.

## Data availability

The X-ray data are deposited into the Protein Data Bank under accession numbers 6MH9(FakB1(A121I)) and 6NM1(FakB1(A158L)). The NMR data are deposited into the Biological Magnetic Resonance Data Bank under the entry IDs 50555 (FakB1) and 50556 (FakB1(A121I)).

## Methods

### Bacterial strains and reagents

Sources of supplies were: Perkin Elmer, [^14^C]16:0 (specific activity, 56.1 mCi/mol); Millipore-Sigma, all FA; Cambridge Isotope Laboratories, Inc., 7,7,8,8- tetradeuteriohexadcanoic acid ([d4]16:0) and algal amino acid mixture (U-^13^C, U-D, U-^15^N); Sigma Aldrich (St. Louis, MO), all reagents for buffers; Fisher Scientific, Casamino acids; New England Biolabs (Ipswich, MA), restriction enzymes; Agilent, QuikChange Lightning Site-Directed Mutagenesis Kit; anti-polyHistidine−alkaline phosphatase antibody (Sigma). High capacity streptavidin agarose beads (ThermoFisher). The bacterial strains and their origins are listed in Table S2. The FakA expression plasmid (pJLB11) and the FakB1 expression plasmid (pCS106) were previously constructed (Parsons et al., 2014). Expression vectors containing FakB1(A121I), FakB1(A158L), FakB1(W180E), FakB1(R205E), and FakB1(R205A) were generated using QuikChange Lightning Site-Directed Mutagenesis Kit and pCS106 as a template (Table S2). FakB1(A121I) For and FakB1(A121I) Rev primers were used to generate the A121I mutant, pPJ583. FakB1(A158L) For and FakB1(A158L) Rev primers were used to generate the A158L mutant, pPJ584 (Table S2). B1 W180E primers 1 and 2 were used to generate the W180E mutant, pPJ593. B1 R205A primers 1 and 2 generated the R205A mutant, pPJ594. B1 R205E primers 1 and 2 generated the R205E mutant, pPJ595 (Table S2). Manufacturer’s protocol was followed for mutagenesis PCR. PCR-products from the mutagenesis underwent Dpn-I digestion and transformation into XL10-Gold ultracompetent cells (carbenicillin 50 μg/ml). Purified plasmids were verified by sequencing. The QuikChange Lightning Site-Directed Mutagenesis Kit was also used to create A121I, A158L and W180E FakB1 mutants in the pCS119 vector used in metabolic labeling experiments. The *fakB1* gene in pCS119 (pB1) was previously created (Gullett et al., 2019) and used as a template. CO A121I F and CO A121I R primers were used to generate the A121I mutant in pCS119 (pA121I). CO A158L F and CO A158L R primers were used to generate the A158L mutant in pCS119 (pA158L). CO B1 W180E Fwd and CO B1 W180E Rev primers were used to generate the W180E mutant in pCS119 (pPJ597) (Table S2).

### Protein expression and purification

Expression plasmids containing *S. aureus* FakA, wild-type FakB1, or FakB1 mutants (A121I, A158L, W180E, R205E, R205A) were transformed into BL21 DE3 *E. coli* electrocompetent cells (Invitrogen) and purified as previously described (Parsons et al., 2014). Briefly, induced proteins were lysed, centrifuged, and the supernatant was incubated at room temperature with 10 mM fatty acid (palmitate for FakB1 and FakB1(A121I), FakB1(W180E), FakB1(R205E), FakB1(R205A), and myristate for FakB1(A158L)) for 1 hr to exchange a known fatty acid into the binding pocket before purification. For triple labeled wild-type FakB1 and FakB1(A121I), colonies from the previously mentioned transformation were grown at 37 °C in minimal media containing carbenicillin (50 mg/ml), glucose (2 g/L), and Casamino acids (2 g/L). Overnight cultures were reinoculated (A_600_= 0.01) into 500 ml fresh media containing glucose (2 g/L) and triple labeled algal amino acids (U-^13^C, U-D, U-^15^N) (2 g/L) instead of Casamino acids. Cultures grew until an A_600_= 0.7 before being induced with IPTG (1 mM) and shaken overnight at 16 °C. Cells were harvested as previously described (Parsons et al., 2014).

### Protein crystallization and structure determination

Crystals of FakB1(A121I) and FakB1(A158L) were grown in 0.1 M MES/imidazole pH 6.5, 0.03 M sodium nitrate, 0.03 M disodium hydrogen phosphate, 0.03 M ammonium sulfate, 12.5% PEG 1000, 12.5% PEG 3350 and 12.5% MPD. These conditions are similar to those of FakB1 and the crystal parameters are also very similar (Cuypers et al., 2019). Crystals were cryoprotected with the mother liquor containing 30% glycerol and flash frozen in liquid nitrogen prior to data collection. Diffraction data (360°) were integrated with XDS (Kabsch, 2010a, b) and scaled using AIMLESS/CCP4 (Evans and Murshudov, 2013). The structures were solved by molecular replacement using MOLREP/CCP4 (Vagin and Isupov, 2001) and the wild-type structure as the search model (PDB:5UTO). In both molecular replacement solutions, residues 163-186 in one of the two molecules in the crystal asymmetric unit were incorrect and were rebuilt into omit maps using ITERATIVE BUILD OMIT MAP/PHENIX (Vagin and Isupov, 2001) The final structures were obtained by cycles of rebuilding and refinement using COOT (Emsley et al., 2010) and PHENIX.REFINE (Afonine et al., 2012). Water molecules were placed both automatically and manually with COOT. In all refinements, 5% of the reflections were excluded for calculation of the R_free_values. The final models were deposited to the PDB database and the molecule and refinement parameters are shown in Table 1.

### NMR sample preparation and data collection

All samples were prepared in an NMR buffer comprising 20 mM Tris, 200 mM NaCl (pH 7.5), 90% H_2_O, 10% D_2_O with 1 mM *S. aureus* wild-type FakB1 or FakB1(A121I). NMR data were collected on Bruker Avance 600 MHz, 700 MHz and 800 MHz spectrometers equipped with TCI triple-resonance cryogenic probes initially at 313 K and later at 293 K. All the relaxation dispersion data were measured at 293 K. Initially, ^15^N, ^13^C double-labeled FakB1 was prepared and standard TROSY based 3D assignment experiments were collected. These included HNCA, HNCACB, CBCA(CO)NH, HNCO, and HN(CA)CO along with 2D TROSY ^1^H-^15^N HSQC. A TROSY-based ^15^N-resolved [^1^H, ^1^H]-NOESY spectrum was also recorded (mixing time of 120 msec). These data allowed 90% of the backbone resonances to be assigned. These assignments were transferred to a 293 K spectrum, and the assignments were confirmed using a 3D HNCA spectrum. A ^2^H, ^15^N, ^13^C triple-labeled FakB1 was also prepared, and TROSY based 3D HNCA, HNCACB, HN(CO)CA spectra were collected along with a TROSY-based ^15^N- resolved [^1^H, ^1^H]- NOESY spectrum (mixing time of 150 msec at 293 K). Assignment of ^2^H, ^15^N, ^13^C labeled FakB1(A121I), was confirmed using a 3D HNCA along with a TROSY-based ^15^N- resolved [^1^H, ^1^H]-NOESY spectrum recorded (mixing time of 150 msec at 293 K at 850 MHz). All the data were processed using BRUKER Topspin version 4.0.5, NMRPipe (v7.9) (Delaglio et al., 1995) and analyzed using CARA (v1.8.4). All spectra were referenced directly using DSS for the ^1^H dimension, ^13^C and ^15^N frequencies were referenced indirectly.

### NMR relaxation dispersion data collection and analysis

^15^N single quantum CPMG (Carr-Purcell-Meiboom-Gill) relaxation dispersion experiments (Loria et al., 1999; Tollinger et al., 2001) were used to study the exchange processes in FakB1 and FakB1(A121I). Relaxation dispersion data sets were recorded on both samples at 293 K using both 600 and 800 MHz spectrometers. Fourteen CPMG frequencies from 50 Hz to 1000 Hz were sampled during a constant-time relaxation interval of 40 ms with an inter-scan delay of 3 s, including a reference spectrum without the constant-time relaxation interval. Amide resonance intensities were converted into effective relaxation rates (Loria et al., 1999; Tollinger et al., 2001) and plotted as a function of the CPMG frequency. Error estimates for R_eff_ were obtained from duplicate measurements at 100, 250 and 500 Hz frequencies as previously described (Korzhnev et al., 2004). The relaxation dispersion data were fit to extract global exchange parameters, including exchange rates and population of the states, along with residue specific values such as ^15^N chemical shift differences between exchanging states and intrinsic ^15^N relaxation rates, R_2,0_. This was accomplished with ShereKhan (Mazur et al., 2013) using Carver-Richards equation (Carver and Richards, 1972) for two-site exchange. Residues that did not overlap with other resonances (Ile59, Thr61, Thr62, Leu165, Lys166, Arg173, Ile174, Val204, Val267, Leu271) were used in the exchange parameter calculation for FakB1. For FakB1(A121I), residues Ile59, Thr61, Thr62, Ser63, Leu165, Arg173, Val204, Gly264, Gly272, and Leu276 were used in the calculation.

### Thermal stability

The structural stabilities of *S. aureus* wild-type FakB1 and mutant derivatives A121I, A158L, W180E, R205E, and R205A were evaluated using thermal shift assays. Five microliters of protein (200 μm) was added to a total of 95 μl of 200 mM MgCl_2_, 150 mM NaCl, 20 mM HEPES (pH 7.5), 2 mM ATP, and Sypro Orange (1:200) (Invitrogen). The plate was centrifuged at 1,500 rpm for 2 min before being placed in an Applied Biosciences 7500 RealTime PCR instrument. A thermal scan from 25 °C to 95 °C was performed using an increment rate of 1 °C/min. The first derivative values of each temperature were calculated and graphed as a function of temperature. All experiments were performed in triplicate.

### Analytical ultracentrifugation

Sedimentation velocity experiments were conducted in a ProteomeLab XL-I analytical ultracentrifuge (Beckman Coulter, Indianapolis, IN) following standard protocols unless mentioned otherwise. Samples in buffer containing 20 mM Tris pH 8, 200 mM NaCl, 10 mM EDTA were loaded into cell assemblies comprised of double sector charcoal-filled centerpieces with a 12 mm path length and sapphire windows. Buffer density and viscosity were determined in a DMA 5000 M density meter and an AMVn automated micro- viscometer (both Anton Paar, Graz, Austria), respectively. The cell assemblies, containing identical sample and reference buffer volumes of 390 μL, were placed in a rotor and temperature equilibrated at rest at 20 °C for 2 h before it was accelerated from 0 to 50,000 rpm. Rayleigh interference optical data as well as absorbance data at 280 nm were collected at 1 min intervals for 12 h. The velocity data were modeled with diffusion-deconvoluted sedimentation coefficient distributions c(s) (Schuck, 2000). The s-value was corrected for time and finite acceleration of the rotor was accounted for in the evaluation of Lamm equation solutions (Zhao et al., 2015). Isotherm data of the signal-average s- values, sw(c), of the total sedimenting system derived from integration of the complete c(s) distributions of all FakA (A) and FakB1 (B) mixtures of sedimentation velocity data at concentrations stated in Table S1 were fitted to a mixed self- and hetero-association model; (A+A) + B + B forming complexes (AA), AB, (AA)B, (AA)BB; self-association of A with two symmetric sites for two B’s. The association scheme used in this analysis was (A+A) + B + B ↔ A +AB + B↔ (AA)B +B↔(AA)(B)2 with the dimer dissociation constant KDAA for the self-association of A and KDAB for the hetero-interaction of A and B with two symmetric sites. Errors of the fits represent the 68.3% confidence interval (CI) using an automated surface projection method (Zhao and Schuck, 2012). Calculations were performed using SEDFIT/SEDPHAT (https://sedfitsedphat.nibib.nih.gov/software/default.aspx). All plots were created in GUSSI (Brautigam, 2015) (http://biophysics.swmed.edu/MBR/software.html).

### FA kinase assays

FA kinase assays contained 0.1 M Tris-HCl (pH 7.5), 10 mM ATP, 20 mM MgCl_2_, 0.2 μm FakA, 20 μM of [^14^C]16:0, Triton X-100 (1%), and fatty acid-free BSA (1 mg/ml). The indicated concentrations of purified FakB proteins were in a total volume of 60 μl. Tubes were incubated at 37 °C for 20 min before acetic acid (0.6%) was added and 40 μl was pipetted on a DE81 Whatman filter paper disc and discs were washed three times, for 20 min each, in ethanol containing acetic acid (1%). Discs were dried and counted by scintillation counting. Apparent Km values were determined performing a nonlinear regression and using the Michaelis-Menten equation. Acyl-phosphate production was also assessed by spotting reactions mixtures on TLC plates. Reactions contained 0.1 M Tris-HCl (pH 7.5), 10 mM ATP (pH 7.5), 20 mM MgCl_2_, either 200 μM liposomes (900 μM DOPG, 10% [^14^C]16:0) or fatty-acid free BSA (1 mg/ml) with [^14^C]16:0 (20 μM), and FakB1 (1 μM for BSA, 0.005 μM for liposomes). The mix was incubated at room temperature for 15 min before FakA (4 μM) was added to begin the reaction. Reagents were incubated at 37 °C for 20 min before acetic acid (0.8%) was added and reactions ceased. Ten microliters of each reaction was spotted on a Silica Gel H plates (Analtech) that were developed with EtOH:CHCl_3_:Et_3_N:H_2_O (34:30:35:6.5) and then imaged on a Typhoon FLA 9500. The bands were quantified using ImageQuant software. Statistical significance was determined using the two-tailed Student’s t test.

### FA incorporation experiments

*S. aureus* strain JLB31 with a plasmid containing either a wild-type *fakB1* gene, a mutant (pA121I, pA158L), or empty plasmid (pCS119) was inoculated into 5 ml of Luria broth containing 0.1% Brij-58, and tubes were shaken at 37 °C until an A_600_ of 0.5 was reached. A FA mixture (10 μM each) of [*d*_4_]16:0, 18:1, and 18:2 was added to the culture and incubated at 37 °C for 30 min before lipids were extracted. All experiments were performed in triplicate and representative spectra are shown. Lipid extracts were resuspended in chloroform:methanol (1:1). PG was analyzed using a Shimadzu Prominence UFLC attached to a QTrap 4500 equipped with a Turbo V ion source (Sciex). Samples were injected onto an Acquity UPLC BEH HILIC, 1.7 μm, 2.1 × 150 mm column (Waters) at 45 °C with a flow rate of 0.2 ml/min. Solvent A was acetonitrile, and solvent B is 15 mM ammonium formate, pH 3. The HPLC program was the following: starting solvent mixture of 96% A / 4% B, 0 to 2 min isocratic with 4% B; 2 to 20 min linear gradient to 80% B; 20 to 23 min isocratic with 80% B; 23 to 25 min linear gradient to 4% B; 25 to 30 min isocratic with 4% B. The QTrap 4500 was operated in the Q1 negative mode. The ion source parameters for Q1 were: ion spray voltage, -4500 V; curtain gas, 25 psi; temperature, 350 °C; ion source gas 1, 40 psi; ion source gas 2, 60 psi; and declustering potential, -40 V. The system was controlled by the Analyst^®^ software (Sciex). The sum of the areas under each peak in the mass spectra was calculated and the percent of each molecular species present was calculated with LipidView software (Sciex). Incorporation of [*d*_4_]16:0 was calculated by combining the values of [*d*_4_]16:0, [*d*_4_]18:0, and [*d*_4_]20:0 molecular species, the values of 18:1 and 20:1 were combined for 18:1 incorporation; the values of 18:2 and 20:2 were combined for 18:2 incorporation. All experiments were performed in triplicate.

### Biotinylated liposome pulldown assay

Five microliters of purified FakB1 (40 μM) was added to 20 µl of high capacity streptavidin agarose beads (ThermoFisher) and 25 μl of liposomes containing the following: 1 mol percent of DOPE-N-(cap biotinyl) and 99 mol percent of DOPC or DOPG; 1 mol percent of DOPE-N-(cap biotinyl), 89 mol percent DOPG, and 10 mol percent of [^14^C]16:0; 1 mol percent of DOPE-N-(cap biotinyl), 79 mol percent DOPG, and 20 mol percent of [^14^C]16:0. Buffer (0.1M Tris, pH 7.5) was added to a total volume of 200 μl and samples were incubated at room temperature on a shaker for 1 hr before being centrifuged at 2,000 x g for 5 min. Supernatant was decanted without disturbing the streptavidin beads, and excess moisture was wicked from the beads before resuspending in an equal volume of buffer. A Western blot was performed on aliquots from the supernatant and pellet using monoclonal anti-polyHistidine−alkaline phosphatase antibody (Sigma). Blots were images on a Typhoon FLA 9500 and band intensities were quantified using ImageQuant (Cytiva). Statistical significance was determined using the two-tailed Student’s t test.

### Surface plasmon resonance (SPR)

SPR experiments were conducted at 25 °C on a Biacore T200 optical biosensor (Cytiva Life Sciences) using the methods of Del Vecchio and Stahelin (Del Vecchio and Stahelin, 2016). A 50:50 mixture of DOPG:DOPC was used for the variable component vesicles, and 100% DOPC was used for the control vesicles allowing for the measurement of net binding to DOPG. Dried lipids (0.5 mM) were hydrated overnight in binding buffer (20 mM Tris pH 7.5, 200 mM NaCl) prior to extrusion. Extruded lipids were captured on an equilibrated, pre-conditioned L1 chip (Cytiva), and the chip was blocked with 3-4 injections of fatty-acid free BSA (0.1 mg/mL). Proteins were prepared in binding buffer as a three-fold dilution series with maximum concentration of 100 μM and injected for 600 s at a flow rate 10 μL/min. The lipid surfaces were regenerated between cycles with NaOH (50 mM) injected for 12 s at flow rate 50 μL/min. The data were processed, double-referenced, and analyzed using the software package Scrubber2 (version 2.0c, BioLogic Software). Equilibrium dissociation constants (*K*_D_) were determined by fitting the data to a 1:1 (Langmuir) equilibrium affinity model.

### Molecular dynamic simulations

Crystal structures of FakB1•16:0 (PDB:5UTO) and FakB1(A121I)•16:0 (PDB:6MH9) were used as initial structures for the simulations. PSFGEN plugin of VMD (Visual Molecular Dynamics) (Humphrey et al., 1996) was used to add a C-terminal carboxylate capping group, an N-terminal ammonium capping group, and hydrogen atoms. The proteins were placed, individually, in a water box using the SOLVATE plugin of VMD. The solvated system was neutralized with Na^+^ and Cl^−^ ions (0.15 M NaCl) using the AUTOIONIZE plugin of VMD. The molecular system was then energy minimized for 2,000 steps and equilibrated for 1 ns. The final equilibrated protein was used for all the consequent membrane-binding simulations. HMMM membranes were employed to capture the membrane-binding conformations of the FakB1•16:0 and FakB1(A121I)•16:0. The short-tailed lipids used in the model enhance lipid diffusion and membrane reorganization thereby allowing spontaneous peripheral protein insertion within the timescales of the simulations (Baylon et al., 2016; Ohkubo et al., 2012; Vermaas et al., 2015). All the independent HMMM membranes were constructed using HMMM BUILDER in CHARMM-GUI (Jo et al., 2008; Qi et al., 2015). Multiple membrane-binding simulations of FakB1•16:0 and FakB1(A121I)•16:0 were performed in the presence of pure PG or PC lipid bilayers and each simulation ran for 100 ns. All membrane-binding simulations began with the protein in the aqueous solution at least 10 Å away from the *cis*-leaflet phosphate plane. The orientation of each protein was varied with respect to the membrane normal (z) axis to ensure the final membrane-bound conformation was not biased by the initial placement.

To test the stability of the membrane-bound conformation of the protein obtained from HMMM simulations (Jo et al., 2008; Qi et al., 2015) a membrane-bound replica of FakB1(A121I)•16:0 was converted to full membrane (full-lipid) using CHARMM-GUI. After another round of equilibration following the CHARMM-GUI protocol, the entire membrane-bound FakB1(A121I)•16:0 was simulated for 500 ns. All simulations were performed in NAMD2 (Phillips et al., 2005; Phillips et al., 2020) using CHARMM36m protein and lipid force fields and TIP3P water (Best et al., 2012; Klauda et al., 2010). Non-bonded interactions were calculated with a 12 Å cutoff and a switching distance of 10 Å. Long-range electrostatic interactions were calculated using the particle mesh Ewald (PME) method (Essmann et al., 1995). The temperature was maintained at 310 K by Langevin dynamics with a damping coefficient of 1.0 ps^-1^. Short tailed HMMM lipids are best simulated in a fixed area ensemble (Ohkubo et al., 2012; Pant and Tajkhorshid, 2020). The pressure was therefore maintained only along the membrane normal (NPnAT) using the Nosé-Hoover Langevin piston method (Martyna et al., 1994).

All analyses were performed in VMD (Humphrey et al., 1996). The membrane-binding configuration and depth of protein insertion into the lipid bilayer were measured by averaging the z positions of all Cα-atoms with respect to the *cis*-leaflet phosphate plane, over the last 50 ns of HMMM simulations. The protein was considered membrane-bound if helix α8’ (residues 177-184) penetrated the bilayer. A heavy-atom cutoff of 3.5 Å was used to define specific lipid-protein contacts. We also monitored the time evolution of the center of mass (COM) of the side-chain heavy atoms of W180, R205, and R209.

