## Supplementary material for "Structural transitions permitting ligand entry and exit in bacterial fatty acid binding proteins"

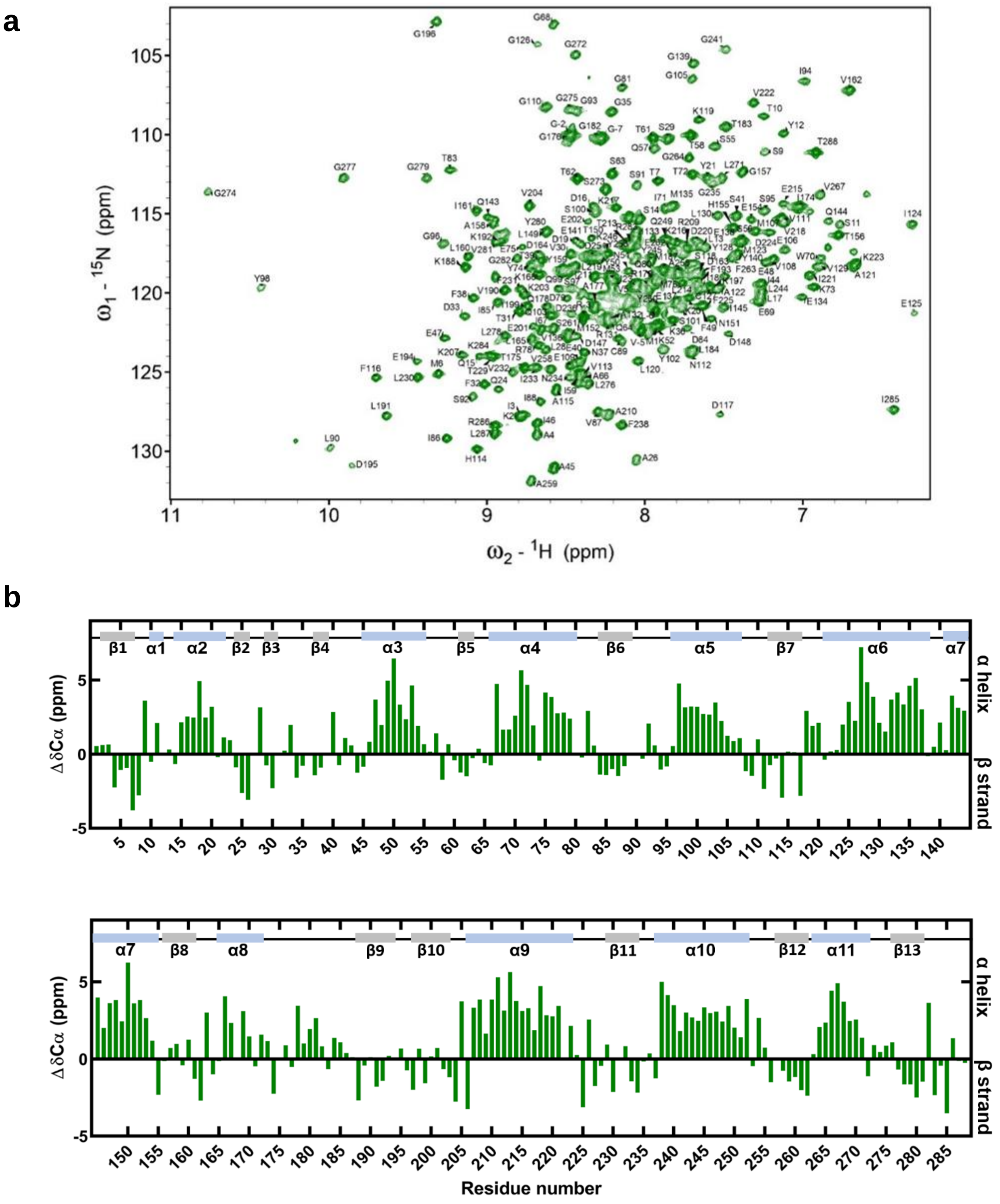


**Supplementary Figure 1. NMR spectroscopy of FakB1.** **a**, Two-dimensional [$^{15}$N, $^{1}$H] TROSY spectra of FakB1 at 293 °K with the assignments as indicated. **b**, Individual residue Cα chemical shift deviation plot at 293 °K for FakB1 correlates with the secondary structure elements (marked on top) observed in the X-ray crystal structure.

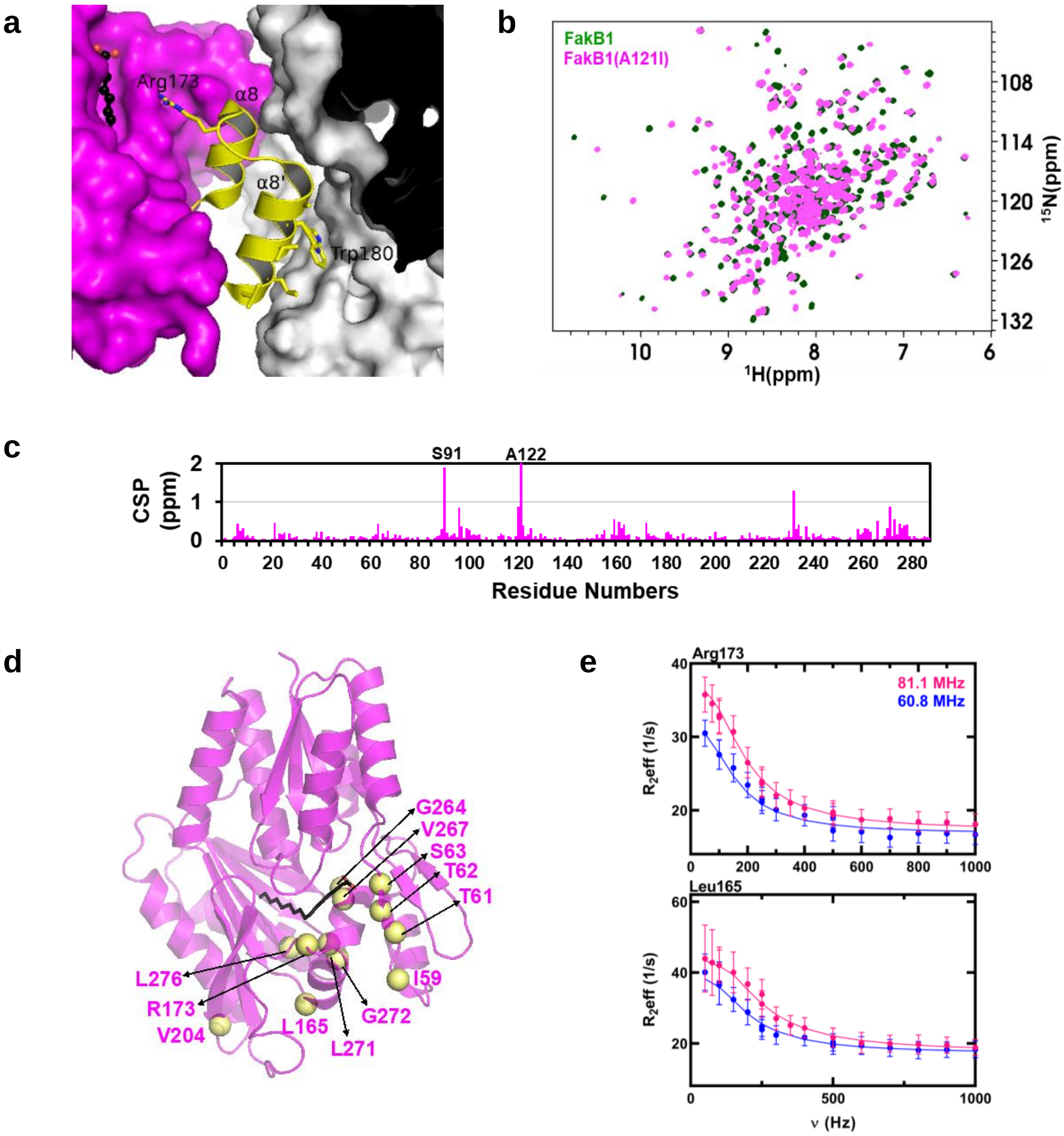


**Supplementary Figure 2. Characterization of FakB1(A121I) and FakB1(A158L).** **a**, View of the α8 and α8′ structure of FakB1(A121I) within the crystal lattice illustrating the packing of α8′ against a hydrophobic patch on an adjacent monomer within the crystal lattice. **b**, Overlay of the TROSY NMR spectra of FakB1 and FakB1(A121I). Asn42, Asn167, Leu168, Ser171, Thr206, His237, Asp240, and Ala268 could not be assigned in the FakB1(A121I) NMR spectrum. **c**, Chemical shift perturbations of FakB1(A121I) with respect to FakB1. **d,** Residues that showed CPMG-RD exchange are indicated as yellow spheres mapped onto the FakB1(A121I) crystal structure. **e**, The two-state global fit (lines) are mapped onto the $^{15}$N relaxation dispersion data at two field strengths for Arg173 and Leu165. Error estimates for Reff were obtained from duplicate measurements at 100, 250, and 500 Hz as described in Methods.

**α4**

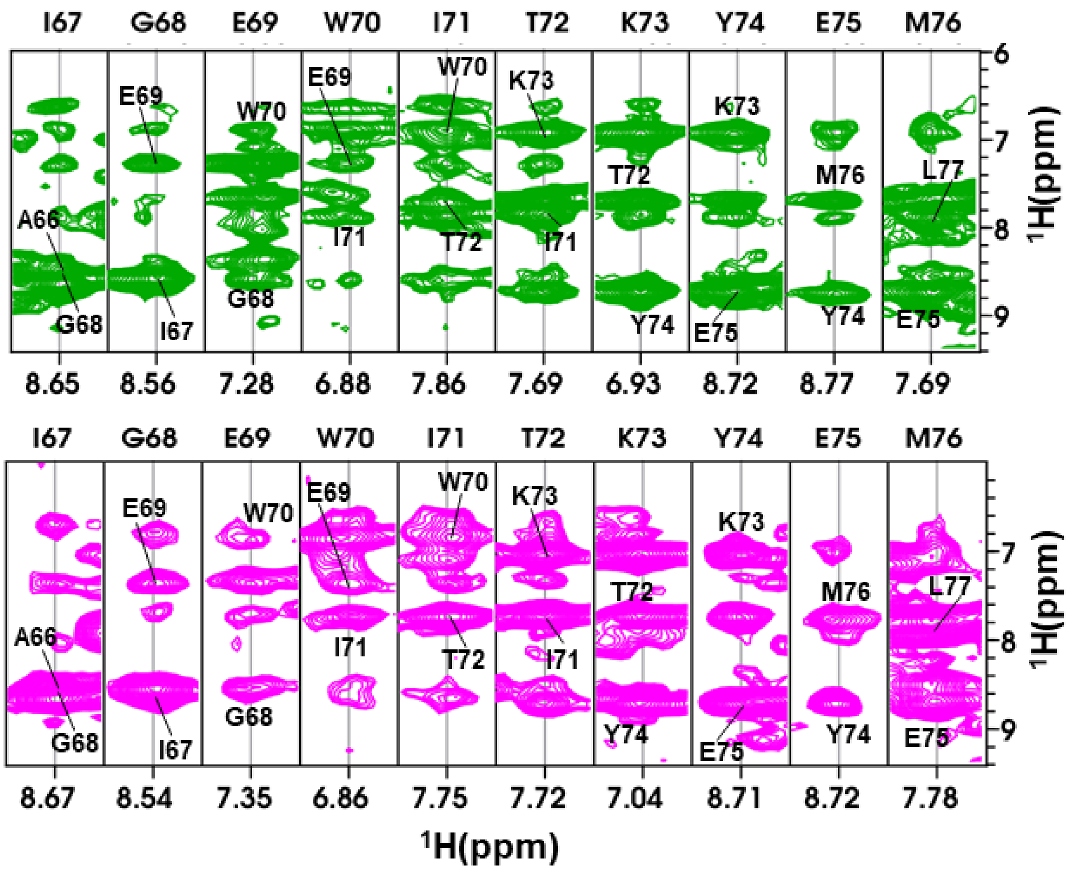


**Supplementary Figure 3. NMR analysis of the FakB1 and FakB1(A121I) solution structure.** NOE contacts between helix α4 in FakB1 (top, green) and FakB1(A121I) (bottom, pink)

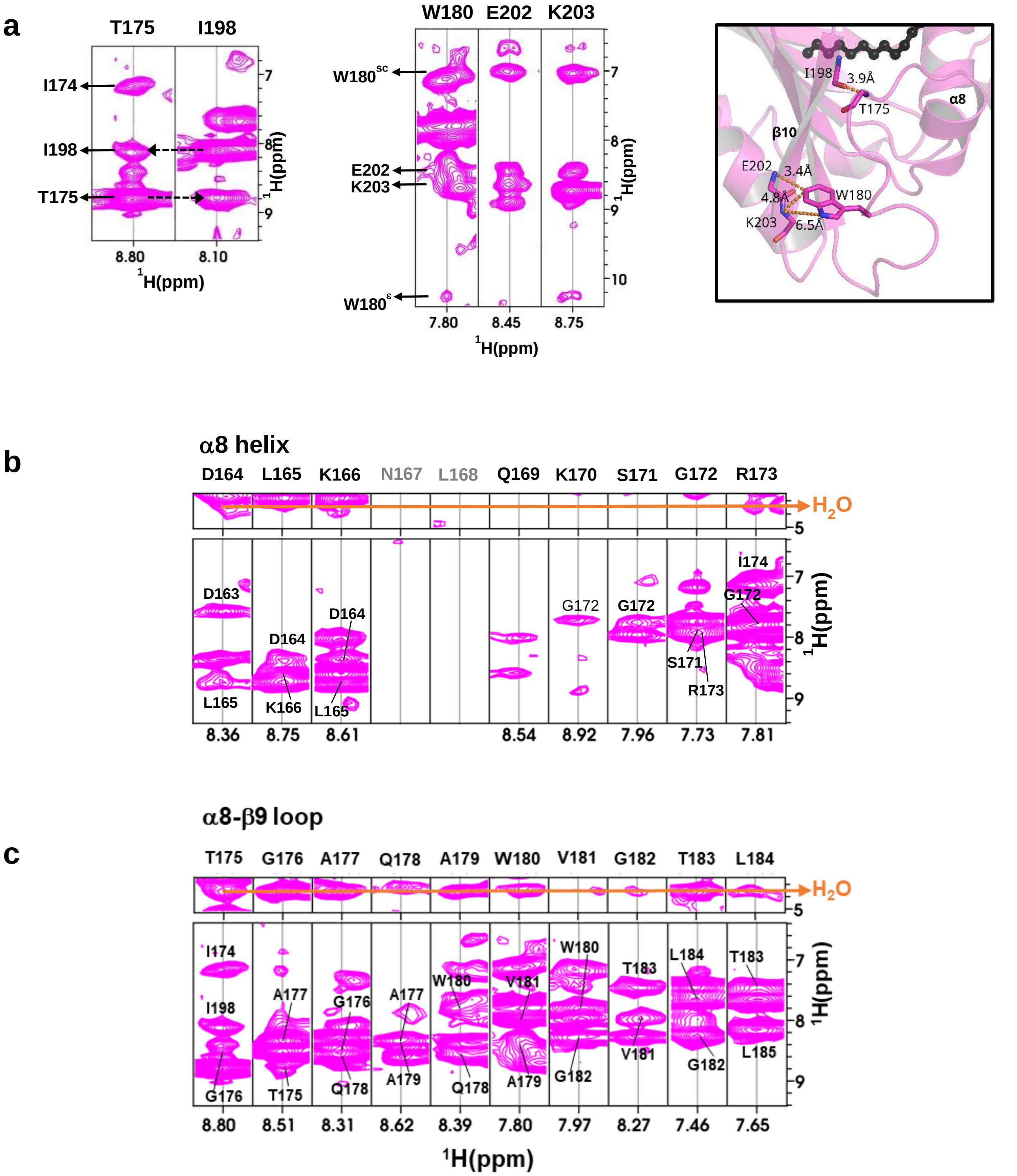


**Supplementary Figure 4. NMR analysis of the FakB1(A121I) solution structure.** **a**, NOE contacts between the α8-β9 loop of domain 2 to residues on domain 1 suggesting the closed conformation shown by the crystal structure. The NOE interactions (dotted orange lines) and computed distances from the NMR data are shown. **b**, Sequential NOEs confirm the existence of helix α8 in solution and the exchange cross peaks with water is shown by the orange line. **c**, The water cross peaks from Thr175-Leu184 are indicative of a structured loop, although the sequential NOEs suggest the partial helical character of the loop centered on Trp180-Val181.

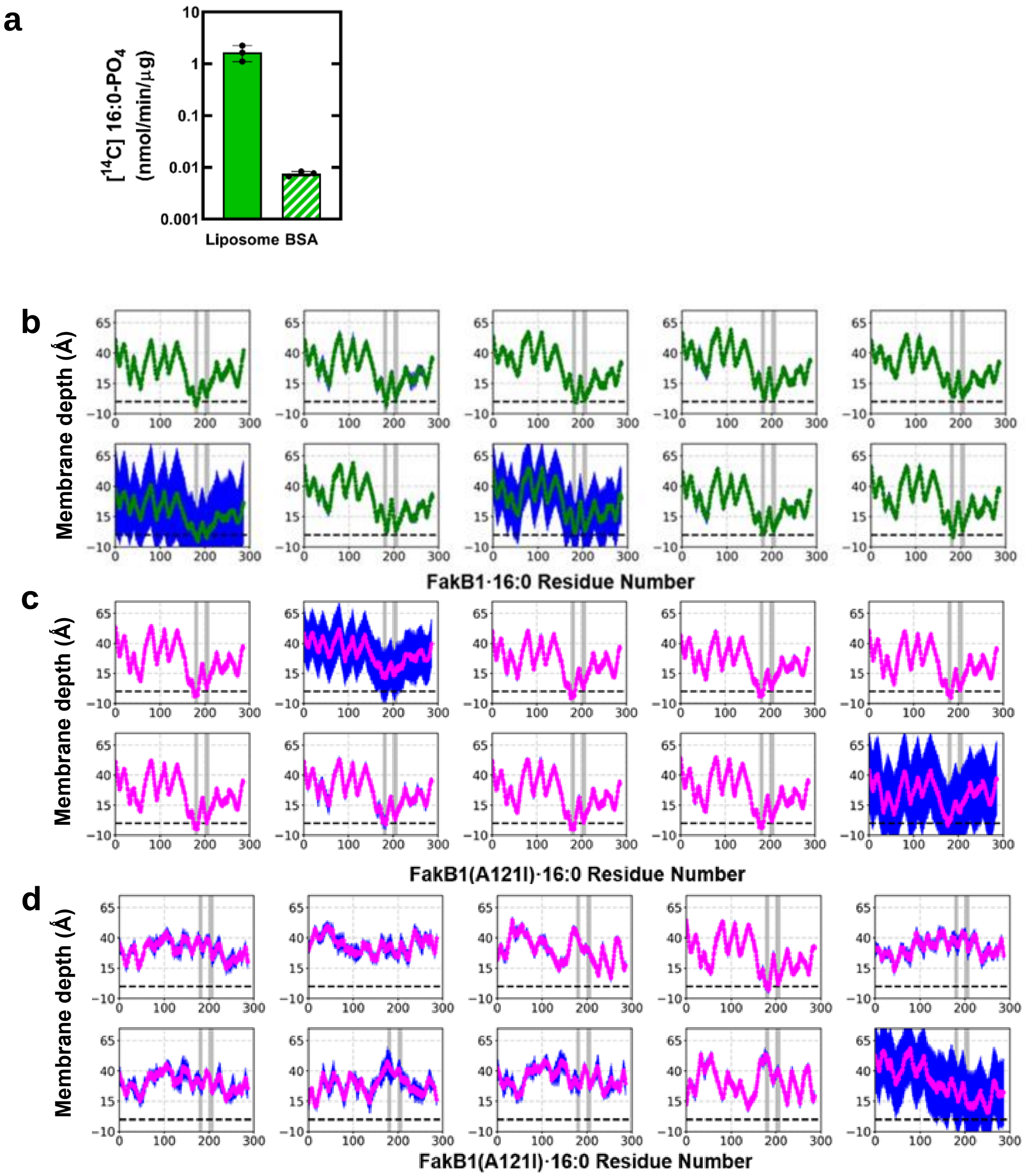


**Supplementary Figure 5. FakB1 exchange assay and the distances of individual FakB1 Cα carbons along the membrane normal calculated during the last 50 ns of HMMM membrane binding simulations.** **a,** FakB1 FA exchange assay using [$^{14}$C]16:0 presented as a BSA-16:0 complex or in PG liposomes. Mean values ± SD are shown; *n* = 3 independent experiments. **b**, Ensemble-averaged Cα distances of FakB1 closed conformation association with PG bilayers. **c**, Ensemble-averaged Cα locations of FakB1(A121I) open conformation binding to PG bilayers. **d**, Ensemble averaged Cα distances for FakB1(A121I) to PC bilayers. In some replicas, we observed high error bars (standard deviations shown in blue) reflecting that the protein is tumbling in solution and membrane binding takes place in last 10 ns. In b-d, dashed lines are used to signify the level of the phosphate layer in the cis monolayer, which was used as a reference (z=0).

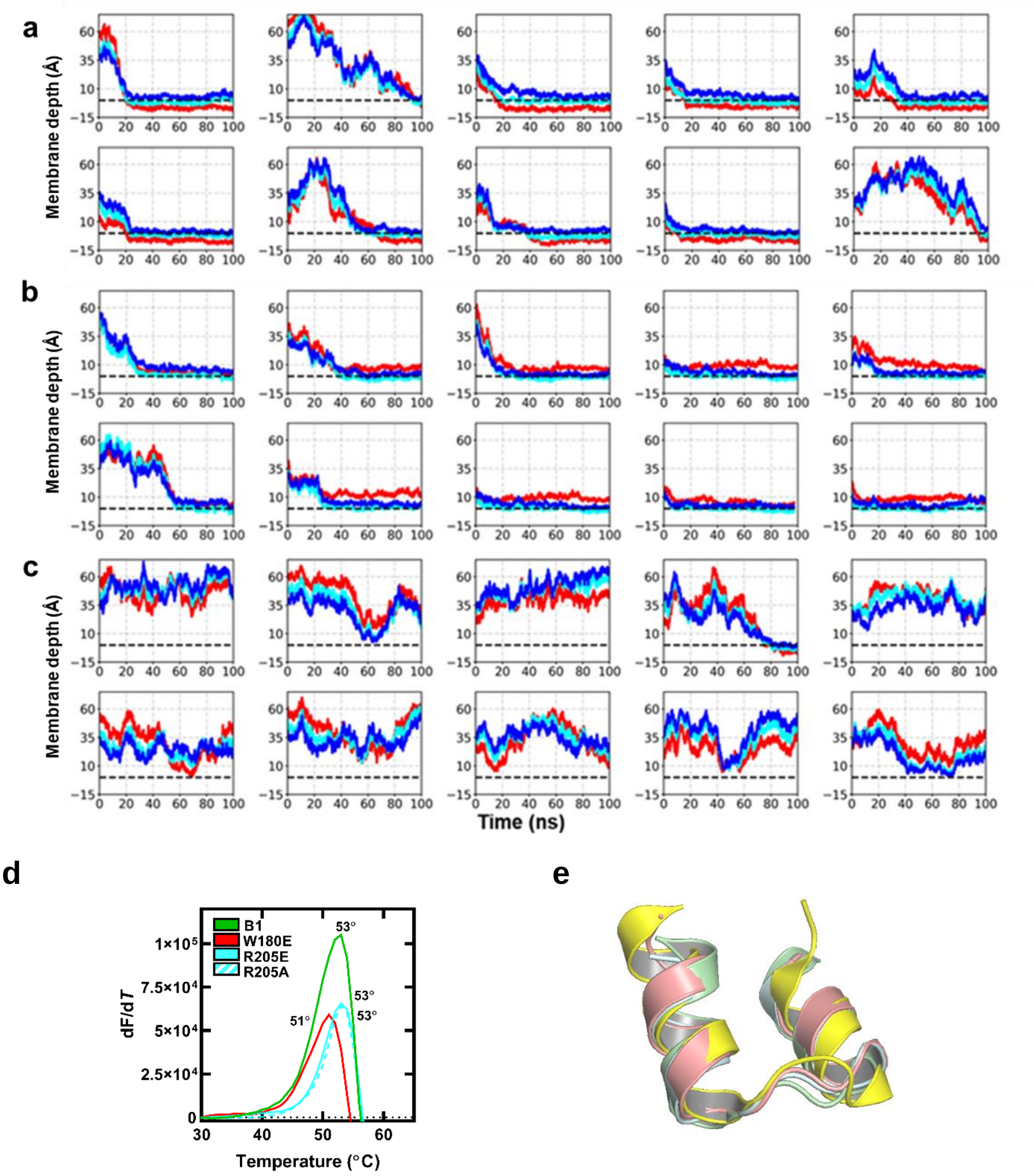


**Supplementary Figure 6. Time evolution of the COM of side chains for three sentinel FakB1 residues along the membrane normal, stability of FakB1 mutants and overlay of FakB1 membrane insertion helices with those in mammalian FABP.** Trp190 (red), Arg205 (cyan) and Arg209 (blue) along the membrane normal (z axis) are plotted as a function of time over 100 ns of 10 independent HMMM membrane binding simulations. **a,** COM distances for membrane insertion of FakB1(A121I) open conformation in PG bilayers. **b**, COM distances for association of FakB1 closed conformation with PG bilayers. **c**, COM distances for FakB1(A121I) in PC bilayers. In a-c, dashed lines are used to signify the level of the phosphate layer in the cis monolayer, which was used as a reference (z=0). **d**, Derivatives of the normalized thermal melting curve using SYPRO orange and 10 µM protein. **e**, Overlay of crystal structures of FakB1(A121I) (PDB: 6MH9, yellow) and human adipocyte FABP4 (PDB: 1TOW, salmon), murine adipocyte FABP4 (PDB: 1A2D, cyan), and rat intestinal FABP2 with palmitate (PDB: 2IFB, green).

**Supplementary Table 1. Sedimentation velocity c(s) analysis of *S. aureus* FakA, FakB1, FakB1(A121I), FakB1(A158L).**

| *Sample* | μM[a] | $s_{20}$ (Svedberg)[b] | $s_{20,w}$ (Svedberg)[c] | Mw (Da) | $f/f_0$[d] |
|---|---|---|---|---|---|
| FakA | 17.84 | 5.81 (78%) | 6.10 | 131,730 | 1.49 |
| FakB1 | 21.62 | 2.74 (86%) | 2.88 | 32,999 | 1.24 |
| FakB1(A121I) | 16.14 | 2.75 (86%) | 2.88 | 33,347 | 1.29 |
| FakB1(A158L) | 17.77 | 2.77 (95%) | 2.91 | 32,740 | 1.24 |
| FakA + | 12.34 | 2.80 (11%) | 2.93 | 48,858 | 1.72 |
| FakB1 | 15.34 | 6.30 (72%) | 6.61 | 182,488 | 1.72 |
| FakA + | 11.72 | 2.58 (9%) | 2.71 | 43,546 | 1.61 |
| FakB1(A121I) | 13.42 | 6.22 (75%) | 6.52 | 162,939 | 1.61 |
| FakA + | 12.48 | 2.62 (10%) | 2.75 | 46,845 | 1.67 |
| FakB1(A158L) | 14.60 | 6.38 (76%) | 6.70 | 177,974 | 1.67 |

[a]Total protein concentrations in μM.

[b]Sedimentation coefficient taken from the ordinate maximum of each peak in the best-fit c(s) distribution at 20°C with percentage protein amount in parenthesis. Sedimentation coefficient (s-value) is a measure of the size and shape of a protein in a solution with a specific density and viscosity at a specific temperature. Values below 5% were not listed.

[c]Standard sedimentation coefficient (s20,w -value) in water at 20°C.

[d]Molar mass values (MW) taken from the c(s) distribution that was transformed to the c(M) distribution. e[f] Best-fit weight-average frictional ratio values $(f/f_0)w$ taken from the c(s) distribution.

**Supplementary Table 2.** Strains, plasmids, and primers used in this study.

| *Strains and Plasmids* | *Description* | *Source* |
|---|---|---|
| **Strains** | | |
| AH1263 | USA300-0114, Erm-sensitive | Boles 2010 |
| JLB31 | *fakB1*:: ΦΝΣ Δ*fakB2* of strain AH1263 | Parson 2014 |
| **Plasmids** | | |
| pCS119 | pCM28SarAP1promoter | Ericson 2017 |
| pB1 | pCS119 expressing *S. aureus* FakB1 | Gullett 2019 |
| pA121I | pCS119 expressing *S. aureus* FakB1(A121I) | This study |
| pA158L | pCS119 expressing *S. aureus* FakB1(A158L) | This study |
| pPJ597 | pCS119 expressing *S. aureus* FakB1(W180E) | This study |
| pET15b | Expression vector | Novagen |
| pET28a | Expression vector | Novagen |
| pJLB11 | *S. aureus* FakA in pET28a | Parson 2014 |
| pCS106 | *S. aureus* FakB1 in pET15b | Parson 2014 |
| pPJ583 | *S. aureus* FakB1(A121I) in pET15b | This study |
| pPJ584 | *S. aureus* FakB1(A158L) in pET15b | This study |
| pPJ593 | *S. aureus* FakB1(W180E) in pET15b | This study |
| pPJ594 | *S. aureus* FakB1(R205A) in pET15b | This study |
| pPJ595 | *S. aureus* FakB1(R205E) in pET15b | This study |
| **Primers** | | |
| FakB1(A121I) For | CTTCGATAGCAAACTGATAGCAATGATTGAAGGTTGC | This study |
| FakB1(A121I) Rev | GCAACCTTCAATCATTGCTATCAGTTTGCTATCGAAG | This study |
| FakB1(A158L) For | GCGTGAACATACCGGTCTCTATCTGATTGTTGATG | This study |
| FakB1(A158L) Rev | CATCAACAATCAGATAGAGACCGGTATGTTCACGC | This study |
| CO A121I F | GTGTTAACGTTCATGCTTTTGATTCTAAACTTATTGCGATGATTGAAGGCT | This study |
| CO A121I R | AGCCTTCAATCATCGCAATAAGTTTAGAATCAAAAGCATGAACGTTAACAC | This study |
| CO A158L F | GATTTAACAAATATGCGTGAACATACAGGCTTATATTTGATTGTTGACGATTTAAAAAATCT | This study |
| CO A158L R | AGATTTTTTAAATCGTCAACAATCAAATATAAGCCTGTATGTTCACGCATATTTGTTAAATC | This study |
| B1 W180E – 1 | GGGTGCCAACCTCTGCCTGTGCACCGGTAATAC | This study |
| B1 W180E – 2 | GTATTACCGGTGCACAGGCAGAGGTTGGCACCC | This study |
| B1 R205E – 1 | CTGAATTGCACGTTTTTTGGTTTCAACTTTTTCTTCCGGGATAATTTTGCCGTC | This study |

| | | |
|---|---|---|
| B1 R205E – 2 | GACGGCAAAATTATCCCGGAAGAAAAAGTTGAAAC CAAAAAACGTGCAATTCAG | This study |
| B1 R205A – 1 | TGCACGTTTTTTGGTAGCAACTTTTTCTTCCGGGAT AATTTTGCC | This study |
| B1 R205A – 2 | GGCAAAATTATCCCGGAAGAAAAAGTTGCTACCAAA AAACGTGCA | This study |
| CO B1 W180E Fwd | AATAACGTACCCACCTCAGCCTGTGCTCCTGTAATT CGG | This study |
| CO B1 W180E Rev | CCGAATTACAGGAGCACAGGCTGAGGTGGGTACGT TATT | This study |
